# Large-scale neural dynamics in a shared low-dimensional state space reflect cognitive and attentional dynamics

**DOI:** 10.1101/2022.11.05.515307

**Authors:** Hayoung Song, Won Mok Shim, Monica D. Rosenberg

## Abstract

Cognition and attention arise from the adaptive coordination of neural systems in response to external and internal demands. The low-dimensional latent subspace that underlies large-scale neural dynamics and the relationships of these dynamics to cognitive and attentional states, however, are unknown. We conducted functional magnetic resonance imaging as human participants performed attention tasks, watched comedy sitcom episodes and an educational documentary, and rested. Whole-brain dynamics traversed a common set of latent states that spanned canonical gradients of functional brain organization, with global synchrony among functional networks modulating state transitions. Neural state dynamics were synchronized across people during engaging movie watching and aligned to narrative event structures. Neural state dynamics reflected attention fluctuations such that different states indicated engaged attention in task and naturalistic contexts whereas a common state indicated attention lapses in both contexts. Together, these results demonstrate that traversals along large-scale gradients of human brain organization reflect cognitive and attentional dynamics.

## Introduction

A central goal in cognitive neuroscience is understanding how cognition arises from the dynamic interplay of neural systems. To understand how interactions occur at the level of large-scale functional systems, studies have characterized neural dynamics as a trajectory in a latent state space where each dimension corresponds to the activity of a functional network (*1–4*). This dynamical systems approach revealed two major insights. First, neural dynamics operate on a low-dimensional manifold. That is, neural dynamics can be captured by a small number of independent latent components due to covariation of neural activity within a system (*5, 6*). Second, neural activity does not just continuously flow along a manifold, but rather systematically transitions between recurring latent “states”, or hidden clusters, within the state space (*7, 8*). Initial work used resting-state neuroimaging (*9–15*) and data simulations (*16–18*) to describe dynamic interactions among brain regions in terms of systematic transitions between brain states.

Less is known about how our mental states—which constantly ebb and flow over time—arise from these brain state transitions. Recent work in human neuroimaging suggests that brain state changes reflect cognitive and attentional state changes in specific contexts (*1, 19*). For example, work has identified neural states during a sustained attention task (*20*) or a working memory task (*21, 22*). Dataset-specific latent states occurred during different task blocks as well as moments of successful and unsuccessful behavioral performance. Another line of work identified latent states during naturalistic movie watching and demonstrated how neural dynamics relate to contents of the movies (*23*) or participants’ ongoing comprehension states (*24*). An open question is whether the *same* latent states underlie cognitive states across all contexts. For example, does the same state underlie successful attention task performance and engaged movie watching? If brain activity traverses a common set of latent states in different contexts, to what extent do the functional roles of these states also generalize?

Shine et al. (*1*) demonstrated that neural dynamics traverse a common low-dimensional manifold across 7 cognitive tasks. The dynamics within this common manifold were aligned to exogenous task blocks and related to individual differences in cognitive traits. Here we expand on this work by probing a common set of latent states that explain neural dynamics during task, rest, and naturalistic contexts in five independent datasets. We also identify the nature of this shared latent manifold by relating it to the canonical gradients of functional brain connectome (*25*). Finally, we relate neural state dynamics to *ongoing* changes in cognitive and attentional states to probe how neural dynamics are adaptively modulated from stimulus-driven and internal state changes.

We collected human fMRI data, the *SitcOm, Nature documentary, Gradual-onset continuous performance task (SONG) neuroimaging dataset*, as 27 participants rested, performed attention tasks, and watched movies. We characterized latent state dynamics that underlie large-scale brain activity in these contexts and related them to cognitive and attentional state changes measured with dense behavioral sampling. Each participant performed 7 fMRI runs over two days: two eye-fixated resting-state runs, two gradual-onset continuous performance task (gradCPT) runs with either face or scene images, two runs of comedy sitcom watching, and a single run of educational documentary watching. The gradCPT measures fluctuations of sustained attention over time (*26, 27*) as participants respond to images (every 1 s) from a frequent category (90% of trials) and inhibit response to images from an infrequent category (10%). The sitcom episodes were the first and second episodes of a South Korean comedy sitcom, *High Kick Through the Roof*, whereas the educational documentary described the geography and history of Korean rivers.

## Functional brain activity transitions between states in a common latent manifold

### Large-scale neural activity transitions between discrete latent states

To infer latent state dynamics, we fit a hidden Markov model (HMM) to probabilistically infer a sequence of discrete latent states from observed fMRI activity (*28*). The observed variables here were the BOLD signal time series from 25 parcels in a whole-brain parcellation of the cortex (17 functional networks) (*29*) and subcortex (8 regions) (*30*) sampled at a 1 s TR resolution (**Fig. 1A** *left*). Parcel time-courses were *z*-normalized within each participant and concatenated across all fMRI runs from all participants. The model inferred two parameters from these time series: the emission probability and the transition probability (see *Methods*). We assumed that the emission probability of the observed variables follows a mixture Gaussian characterized by the mean and covariance of the 25 parcels in each latent state (**Fig. 1A** *right*). The inferred parameters of the model were used to decode latent state sequences. Four was chosen as the number of latent states (*K* = 4) based on the optimal model fit to the data when tested with leave-one-subject-out cross-validation (chosen among *K* of 2 to 10; **fig. S1**).

**Fig. 1.**
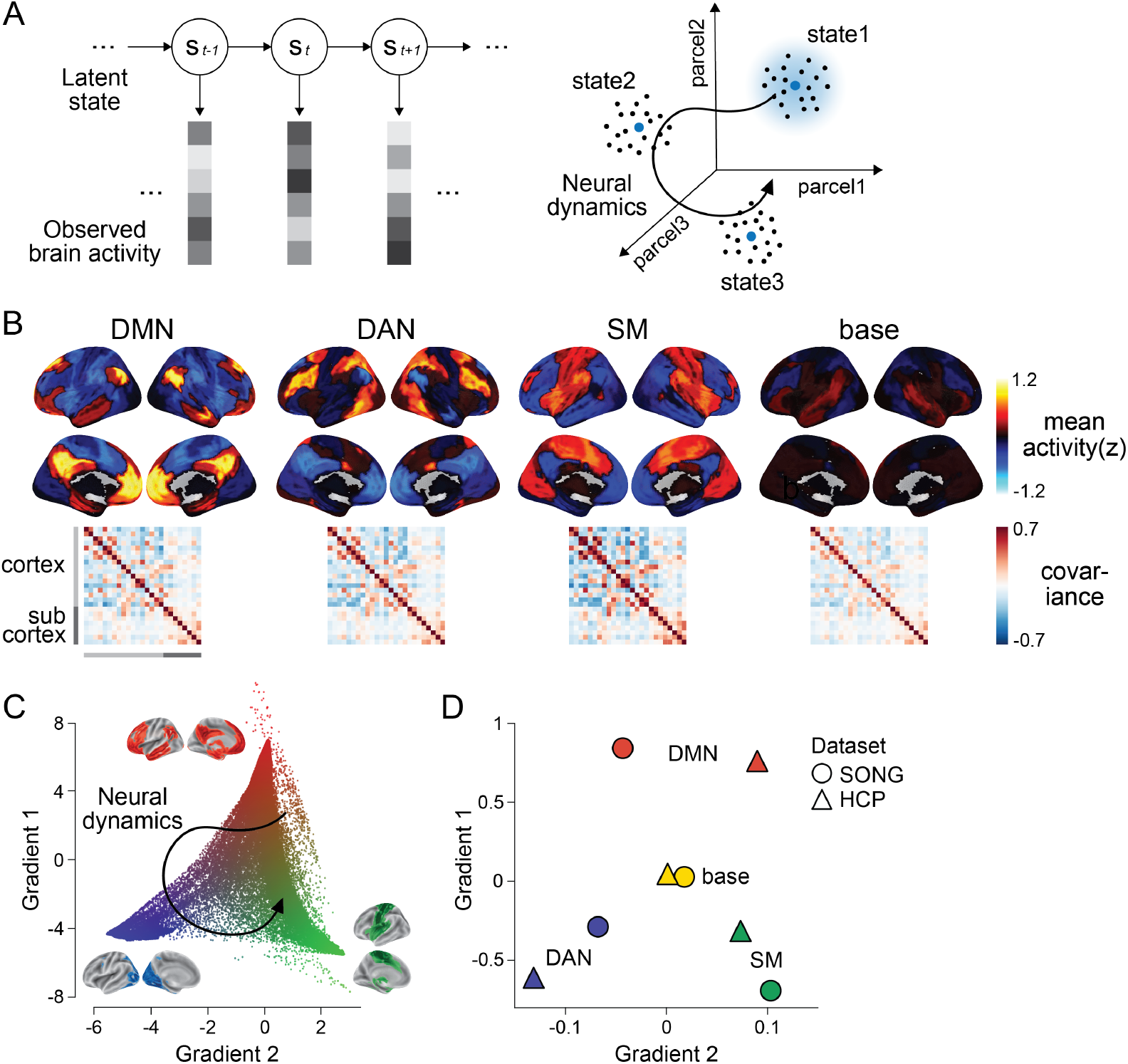
Latent state space of the large-scale neural dynamics. **(A)** Schematic illustration of the HMM inference. (Left) The HMM infers a discrete latent state sequence from the observed 25-parcel fMRI time series. (Right) The fMRI time course can be visualized as a trajectory within a 25-dimensional space, where black dots indicate activity at each moment in time. The HMM probabilistically infers discrete latent clusters within the space, such that each state can be characterized by the mean activity (blue dots) and covariance (blue shaded area) of the 25 parcels. The figure is adapted from Cornblath et al. (*22*). **(B)** Four latent states inferred by the HMM fits to the SONG dataset. Mean activity (top) and pairwise covariance (bottom) of the 25 parcels’ time series is shown for each state. See **fig. S6** for replication with the Human Connectome Project [HCP] dataset. **(C)** Conceptualizing low-dimensional gradients of functional brain connectome as a latent manifold of large-scale neural dynamics. Each dot corresponds to a cortical or subcortical voxel situated in gradient space. The colors of the brain surfaces (inset) indicate voxels with positive or negative gradient values with respect to the nearby axes. Data and visualizations are adopted from Margulies et al. (*25*). **(D)** Latent neural states situated in gradient space. Positions in space reflect the mean element-wise product of the gradient values of the 25 parcels and mean activity patterns of each HMM state inferred from the SONG (circles) and HCP (triangles) datasets.

**Figure 1B** illustrates the four latent neural states inferred by the HMM in the SONG dataset (see **fig. S2** for condition-specific latent states). We labeled three states the default mode network (DMN), dorsal attention network (DAN), and somatosensory motor (SM) states based on high activation of these canonical brain networks (*29*). (Note that these state labels are only applied for convenience. Each state is characterized by whole-brain *patterns* of activation, deactivation, and covariance, rather than simply corresponding to activation of the named network.) The fourth state was labeled the ‘base’ state because activity was close to baseline (*z* = 0) and covariance strength (i.e., the sum of the absolute covariance weights of the edges) was comparatively low during this state. The SM state, on the other hand, exhibited the highest covariance strength, whereas the covariance strengths of the DMN and DAN states were comparable. Compared to null latent states derived from surrogate fMRI time series, the four states exhibited activity patterns more similar to large-scale functional systems (*31–34*) and significantly higher covariance strength (see **fig. S3** for examples of null latent states). These states were replicated with 250 regions of interest (ROIs) consisting of 200 cortical (*35*) and 50 subcortical regions (*30*), albeit with a caveat that the HMM provides a poorer fit to the higher-dimensional time series (**fig. S4**). Neural state inference was robust to the choice of K (**fig. S1**) and the fMRI preprocessing pipeline (**fig. S5**).

To validate that these states are not just specific to the SONG dataset, we analyzed fMRI data from the Human Connectome Project (HCP; N=119) (*36*) collected during rest, 7 block-designed tasks—the emotion processing, gambling, language, motor, relational processing, social cognition, and working memory tasks (*37*)—and movie watching (*38*). The same HMM inference was conducted independently on the HCP dataset using *K* = 4 (**fig. S6**). HCP states closely mirrored the DMN, DAN, SM, and base states (Pearson’s correlations between activity patterns of SONG- and HCP-defined states: DMN: 0.831, DAN: 0.814, SM: 0.865, base: 0.399). Thus, the latent states are reliable and generalize across independent datasets.

### Latent state dynamics span low-dimensional gradients of the functional brain connectome

HMM results demonstrate that large-scale neural dynamics in diverse cognitive contexts (tasks, rest, and movie watching) are captured by a small number of latent states. Intriguingly, the DMN, DAN, and SM systems that contribute to these states tile the principal gradients of large-scale functional organization. In a seminal paper, Margulies et al. (*25*) applied a nonlinear dimensionality reduction algorithm to capture the main axes of variance in the resting-state static functional connectome of 820 individuals. They found that the primary gradient dissociated unimodal (visual and SM regions) from transmodal (DMN) systems. The secondary gradient fell within the unimodal end of the primary gradient, dissociating the visual processing from the SM systems. These gradients, argued to be an “intrinsic coordinate system” of the human brain (*39*), reflect variations in brain structure (*40–42*), gene expressions (*43*), and information processing (*39*).

We hypothesized that the spatial gradients reported by Margulies et al. (*25*) act as a low-dimensional manifold over which large-scale dynamics operate (*44*), such that traversals within this manifold explain large variance in neural dynamics and, consequently, cognition and behavior (**Fig. 1C**). To test this idea, we situated the mean activity values of the four latent states along the gradients defined by Margulies et al. (*25*) (see *Methods*). The DMN, DAN, and SM states fell at the ends of the primary and secondary gradients, whereas the base state was located at the center of the gradient space (**Fig. 1D**).

We asked whether these predefined gradients explain as much variance in neural dynamics as the latent subspace optimized for a specific dataset. To do so, we applied principal component analysis to the 25-parcel time series of the SONG and HCP datasets (**fig. S7**). The top principal components identified from each dataset closely resembled the predefined gradients, with significant Pearson’s correlations observed for the first (SONG: 0.896, FDR-*p* = 0.001; HCP: 0.897, FDR-*p* = 0.003) and second (SONG: 0.672, FDR-*p* = 0.001; HCP: 0.330, FDR-*p* = 0.394) components. Although, by definition, the gradients explained less variance in the parcel time courses than the principal components (paired *t*-tests, 1st: SONG: *t*(187) = 12.862, HCP: *t*(3092) = 50.135, FDR-*p* values < 0.001; 2nd: *t*(187) = 12.990, *t*(3092) = 13.318, FDR-*p* values < 0.001), differences were minor (*r*^2^ of 0.007–0.015 difference) and the gradients explained significant variance (1st: SONG: *r*^2^ = 0.169, HCP: 0.155; 2nd: SONG: 0.110, HCP: 0.120; FDR-*p* values < 0.001). The striking similarity between the dataset-specific principal components and predefined gradients, with comparable explained variance, suggest that the low-dimensional state space of the whole-brain *dynamics* closely recapitulates gradients defined by the *static* functional brain connectome. This suggests that the spatial gradients may operate as a shared low-dimensional manifold of large-scale neural dynamics that can generalize across contexts and datasets (*44*).

### Transient global desynchrony precedes neural state transitions

Neural states constantly change over time. State transitions can be construed as traversals in a low-dimensional space whose axes are defined by principal gradients of functional brain organization. When and how do these neural state transitions occur? What indicates that the system is likely to transition from one state to another?

We predicted that neural state transitions are related to changes in interactions between functional networks. To test this account, we computed cofluctuation between all pairs of parcels at every TR (1 s). Cofluctuation operationalizes the time-resolved interaction of two regions as an absolute element-wise product of their activity at every time step after *z*-normalization of their time series (*45–47*). We time-aligned cofluctuation values to moments of neural state transitions estimated from the HMM (**Fig. 2A**). A decrease in cofluctuation prior to the neural state transitions (at time *t-1*) was observed for every pair of cortico-cortical networks (*z* = 645.75, FDR-*p* = 0.001). Cortico-subcortical pairs (*z* = 424.05, FDR-*p* = 0.001) and subcortico-subcortical connections (*z* = 64.85, FDR-*p* = 0.037) also showed decreased cofluctuation before state transitions, although the effects were less pronounced, especially for subcortico-subcortical connections (paired Wilcoxon signed rank tests comparing the degrees of cofluctuation change, FDR-*p* values < 0.001). Results were replicated with the 250-ROI parcellation scheme as well as with the HCP dataset (**fig. S8**). Furthermore, repeating this analysis with null HMMs on circular-shifted time series suggests that the effect is not simply a byproduct of the chosen computational model (**fig. S9**). These results are consistent with prior empirical findings that desynchronization, a “transient excursion away from the synchronized manifold” (*48*), allows the brain to switch flexibly between states (*18, 49–51*).

**Fig. 2.**
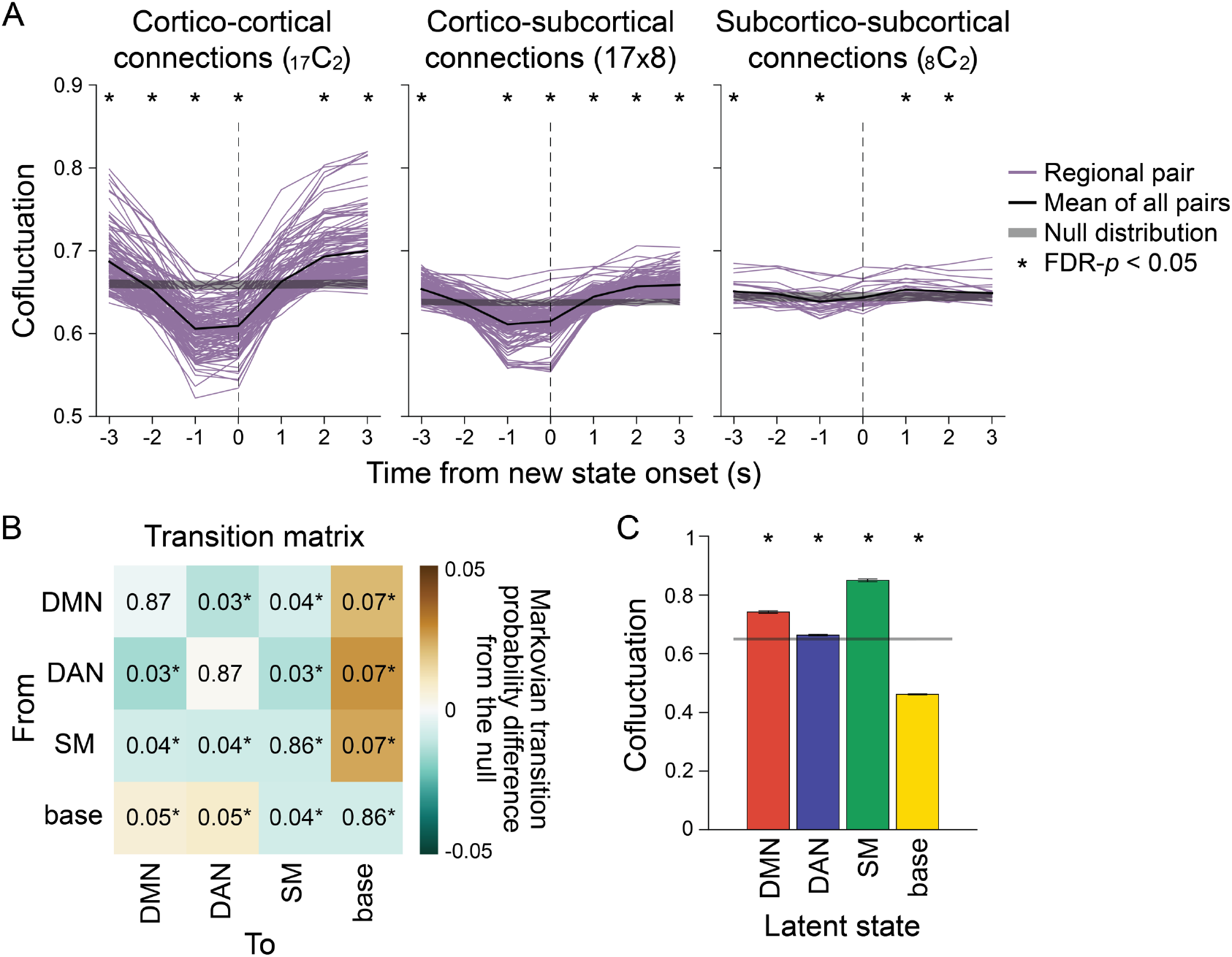
Neural state transitions. **(A)** Changes in cofluctuation of the parcel pairs, time-aligned to HMM-derived neural state transitions. State transitions occur between time *t*-1 and *t*. Purple lines indicate the mean cofluctuation of cortico-cortical (left), cortico-subcortical (middle), and subcortico-subcortical (right) parcel pairs across fMRI runs and participants, and the thick black line indicates the mean of these pairs. The shaded gray area indicates the range of the null distribution (mean ± 1.96 × standard deviation), in which the moments of state transitions were randomly shuffled (asterisks indicate FDR-*p* < 0.05). **(B)** Transition matrix indicating the first-order Markovian transition probability from one state (row) to the next (column), averaged across all participants’ all fMRI runs. The values indicate transition probabilities, such that values in each row sums to 1. The colors indicate differences from the mean of the null distribution where the HMMs were conducted on the circular-shifted time series. **(C)** Mean degrees of global cofluctuation at moments of latent neural state occurrence. The measurements at each time point were averaged within participant based on latent state identification, and then averaged across participants. The bar graph indicates the mean of all participants’ all fMRI runs. The error bars indicate S.E.M. The shaded gray area indicates the range of the null distribution, in which the analyses were conducted on the circular-shifted latent state sequence. See **fig. S8** for replication with the HCP dataset.

### The base state acts as a flexible hub in neural state transitions

To further address how neural state transitions occur, we analyzed the HMM’s transition matrix, which indicates the probability of a state at time *t-1* transitioning to another state or remaining the same at time *t*. The probability of remaining in the same state was dominant (>85%), whereas the probability of transitioning to a different state was less than 8% (**Fig. 2B**; **fig. S10**). To investigate whether certain state transitions occurred more often than expected by chance, we compared the transition matrix to a null distribution where the HMM was conducted on circular-shifted fMRI time series. The DMN, DAN, and SM states were more likely to transition to and from the base state and less likely to transition to and from one another than would be expected by chance (**Fig. 2B, fig. S10**; FDR-*p* values < 0.05). The result suggests that the base state acts as a hub in neural state transitions.

Given that global desynchrony indicates moments of neural state transitions (**Fig. 2A**), we used this measure to validate the role of the base state as a “transition-prone” state. Cofluctuation between every pair of parcels was computed at every TR, which was averaged across parcel pairs to represent a time-resolved measure of global cofluctuation (**Fig. 2C**). When comparing the degree of global cofluctuation across the four latent states, we found that the base state exhibited the lowest degree of global cofluctuation (paired *t*-tests comparing cofluctuation in base state vs. DMN, DAN, and SM states, SONG: *t*(187) > 61, HCP: *t*(3093) > 170, FDR-*p* values < 0.001) which was significantly below chance (FDR-*p* values < 0.001). This suggests that the base state was the most desynchronized state among the four, potentially operating as a transition-prone state. Low global synchrony during the base state was not driven by spurious head motion (**fig. S11**). Thus, the base state, situated at the center of the gradient space, is a flexible “hub” state with a high degree of freedom to transition to other functionally specialized states.

## Neural state dynamics are modulated by ongoing cognitive and attentional states

### Latent state dynamics differ across contexts and are synchronized during movie watching

We identified four latent states that are common across rest, task performance, and movie watching. Although the latent manifold of neural trajectories may be shared across contexts, different subspaces may be occupied by different contexts. For example, one state may occur in one context but not in others. We asked whether the pattern with which brain activity “visits” the four states differed across contexts.

We used the HMM to infer the latent state sequence of each fMRI run (**Fig. 3A**) and summarized the fractional occupancy of each state (i.e., proportion of time that a state occurred) (**Fig. 3B**). All four states occurred in all fMRI runs, with no state occurring on more than 50% of time points in a run. Thus, these states are common across contexts rather than specific to one context. Fractional occupancy, however, differed across rest, task, and naturalistic contexts, with strikingly similar values between runs of similar contexts (e.g., rest runs 1 and 2). In contrast to the similar fractional occupancy values of the two sitcom-episode runs, fractional occupancy in the documentary watching condition differed despite the fact that it also involved watching an audiovisual stimulus. During the documentary, the base state occurred less frequently whereas the SM state occurred more frequently than during the sitcom episodes.

**Fig. 3.**
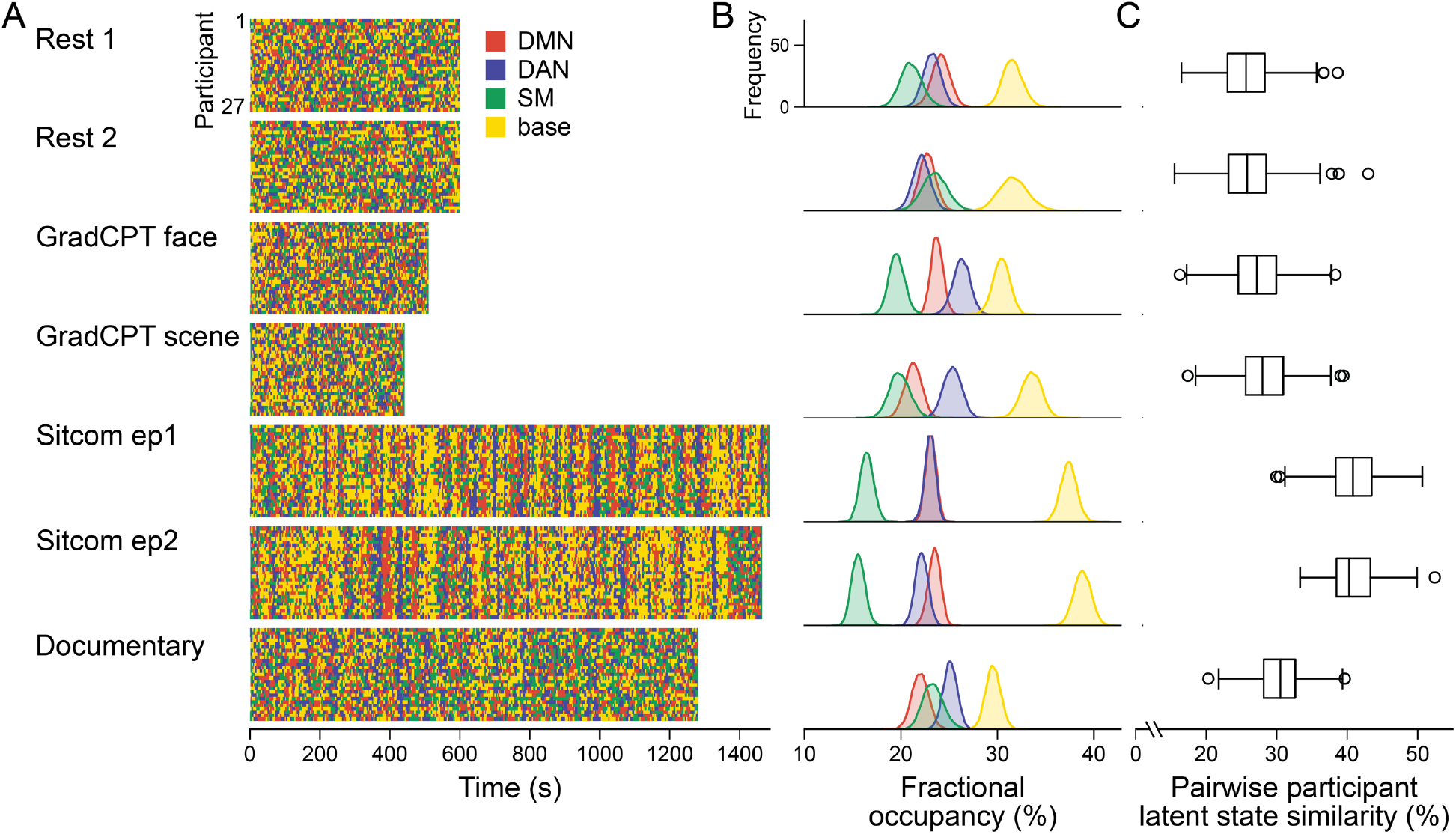
Latent neural state dynamics in the seven fMRI runs. **(A)** Latent state dynamics inferred by the HMM for all participants. Colors indicate the state that occurred at each time point. **(B)** Fractional occupancy of the neural states in each run. Fractional occupancy was calculated for each individual as the ratio of the number of time points at which a neural state occurred over the total number of time points in the run. Distributions indicate bootstrapped mean of the fractional occupancies of all participants. The chance level is at 25%. **(C)** Synchrony of latent state sequences across participants. For each pair of participants, sequence similarity was calculated as the ratio of the number of time points when the neural state was the same over the total number of time points in the run. Box and whisker plots show the median, quartiles, and range of the similarity distribution.

Latent state dynamics were synchronized across participants watching the comedy sitcom episodes (mean pairwise participant similarity: episode 1: 40.81 ± 3.84%, FDR-*p* = 0.001; episode 2: 40.79 ± 3.27%, FDR-*p* = 0.001; paired comparisons, non-parametric *p* = 0.063; **Fig. 3C**). Less synchrony was observed between participants watching the educational documentary (30.39 ± 3.38 %, FDR-*p* = 0.001; paired comparisons with the two sitcom episodes, both *p* < 0.001). No significant synchrony was observed during the resting-state runs (run 1: 25.81 ± 4.00 %, FDR-*p* = 0.230; run 2: 25.84 ± 4.08 %, FDR-*p* = 0.183).

These results were replicated when we applied the SONG-trained HMM to decode latent sequences of the three independent datasets (**fig. S12**). The four neural states occurred in every run of every dataset tested, with maximal fractional occupancies all below 50%. Intersubject synchrony of the latent state sequence was high during movie watching and story listening but at chance during rest. Together the results validate that neural states identified from the SONG dataset generalize not only across contexts but also to independent datasets.

Prior studies reported that regional activity (*52, 53*) and functional connectivity (*54–56*) are synchronized across individuals during movie watching and story listening, and that attentional engagement modulates the degree of intersubject synchrony (*57–59*). Our results indicate that the intersubject synchrony occurs not only at regional and pairwise regional scales, but also at a global scale via interactions of functional networks. Furthermore, stronger entrainment to the stimulus during sitcom episodes compared to documentary watching condition suggests that overall attentional engagement may mediate the degree of large-scale synchrony (mean reports on overall engagement from a scale of 1 [not at all engaging] to 9 [completely engaging]: sitcom episode1: 6.78 ± 1.05, episode2: 6.93 ± 1.41, documentary: 3.59 ± 1.21). Indeed, demonstrating a relationship between neural state dynamics and narrative engagement (**Supplementary Text**), participant pairs that exhibited similar engagement dynamics showed similar neural state dynamics (sitcom episode 1: Spearman’s *r* = 0.274, FDR-*p* = 0.005; episode 2: *r* = 0.229, FDR-*p* = 0.010; documentary: *r* = 0.225, FDR-*p* = 0.005).

### Neural state dynamics are modulated by narrative event boundaries

Latent state dynamics are synchronized across individuals watching television episodes and listening to stories, which suggests that latent neural states are associated with shared cognitive states elicited by an external stimulus. How are these neural state dynamics modulated by stimulus-driven changes in cognition?

Our comedy sitcom episodes had unique event structures. Scenes alternated between two distinct storylines (A and B) that took place in different places with different characters. Each episode included 13 events (7 events of story A and 6 events of B) ordered in an ABAB sequence. This structure required participants to switch between the two storylines at event boundaries and integrate them in memory to form a coherent narrative representation (*60–62*).

We asked if any latent state consistently occurred at narrative event boundaries (**Fig. 4A**). In both sitcom episodes, the DMN state was more likely to occur than would be expected by chance after event boundaries (∼50% probability, FDR-*p* < 0.01), complementing past work that showed the involvement of the DMN at event boundaries (*63–65*). The base state, on the other hand, was less likely to occur after event boundaries (∼10% probability). DAN and SM state occurrences were not modulated by event boundaries (**fig. S13**). These results replicated when the SONG-defined HMM was applied to a 50-min story-listening dataset (*66*) in which 45 events were interleaved in an ABAB sequence (**Fig. 4B**). A transient increase in hippocampal BOLD activity occurred after event boundaries (**fig. S13**), replicating previous work (*63, 65, 67, 68*). Together, our results suggest that event boundaries affect neural activity not only at a regional level, but also at a whole-brain systems level.

**Fig. 4.**
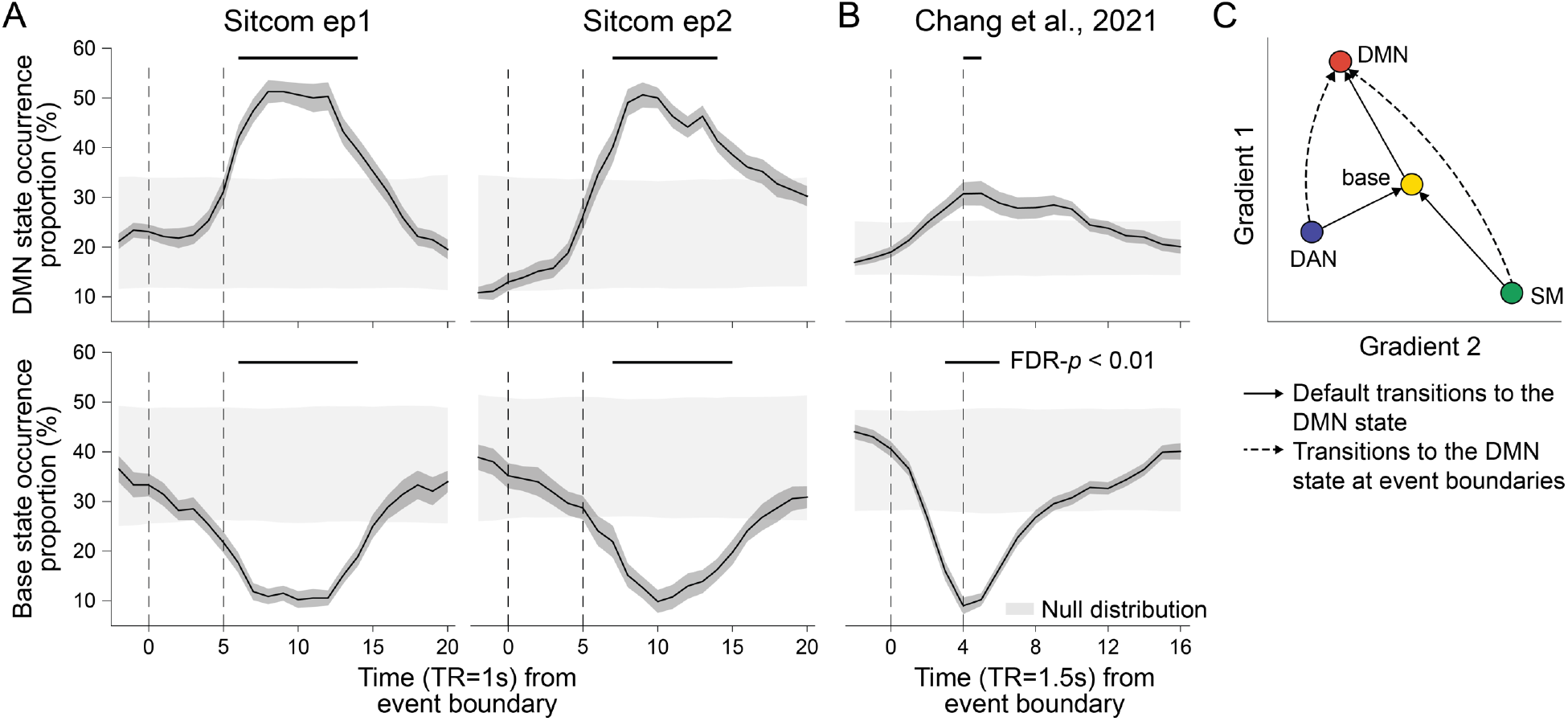
Neural state occurrence and transitions at narrative event boundaries. **(A)** The proportion of the DMN (top) and base state (bottom) occurrences time-aligned to narrative event boundaries of sitcom episode 1 (left) and 2 (right). State occurrence at time points relative to the event boundaries per stimulus was computed within participant and then averaged across participants. The dark gray shaded areas around the thick black line indicate S.E.M. The dashed lines at *t*=0 indicate moments of new event onset and the lines at *t*=5 account for hemodynamic response delay of the fMRI. The light gray shaded areas show the range of the null distribution in which boundary indices were circular-shifted (mean ± 1.96 × standard deviation), and the black lines on top of the graphs indicate statistically significant moments compared to chance (FDR-*p* < 0.01). **(B)** The proportion of the DMN (top) and base state (bottom) occurrence time-aligned to narrative event boundaries of Chang et al.’s (*66*) audio narrative. Latent state dynamics were inferred based on the HMM trained on the SONG dataset. Lines at *t*=4 account for hemodynamic response delay. **(C)** Schematic transitions to the DMN state at narrative event boundaries (dashed lines), compared to the normal trajectory which pass through the base state (solid lines). See **fig. S14** for results of statistical analysis.

How does brain activity transition to the DMN state at event boundaries? To investigate how event boundaries perturb neural dynamics, we compared transitions to the DMN state that occurred at event boundaries (i.e., between 5 and 15 s after boundaries) to those that occurred at the rest of the moments (non-event boundaries) (**fig. S14**). At non-event boundaries, the DMN state was most likely to transition from the base state, accounting for more than 50% of the transitions to the DMN state. Interestingly, however, at event boundaries, base-to-DMN state transitions significantly dropped while DAN-to-DMN and SM-to-DMN state transitions increased (**Fig. 4C**). A repeated-measures ANOVA showed a significant interaction between the latent states and the event boundary conditions (sitcom episode 1: F(2,50) = 10.398; episode 2: F(2,52) = 12.794; Chang et al.: F(2,48) = 31.194; all *p* values < 0.001). Thus, although the base state typically acts as a transitional hub (**Fig. 2B**), neural state transitions at event boundaries are more likely to occur directly from the DAN or SM state to the DMN state without passing through the base state due to the DMN state’s functional role at event boundaries. These results illustrate one way in which neural systems adaptively reconfigure in response to environmental demands.

### Neural state dynamics reflect attention dynamics in task and naturalistic contexts

In addition to changes in cognitive states, sustained attention fluctuates constantly over time (*26, 69–73*). Previous studies showed that large-scale neural dynamics that evolve over tens of seconds capture meaningful variance in attentional states (*20, 73–75*). We asked whether latent neural state dynamics reflect ongoing changes in attention. To infer participants’ attentional fluctuations during the gradCPT, we recorded response times (RT) to every frequent-category trial (∼1 s). The RT variability time course was used as a proxy for fluctuating attentional state, with moments of less variable RTs (i.e., stable performance) indicating attentive states (**Fig. 5A** and **B**). Paying attention to a comedy sitcom, on the other hand, involves less cognitive effort than attending to controlled psychological tasks (*76–79*) and is affected by narrative-evoked cognitive processes such as emotion (*80, 81*), social cognition (*82, 83*) or causal reasoning (*24, 84*). To assess participants’ fluctuating levels of attentional engagement during the sitcom episodes and documentary, we asked participants to continuously self-report their levels of engagement on a scale of 1 (Not engaging at all) to 9 (Completely engaging) as they re-watched the stimuli outside the fMRI (**Fig. 5A** and **B**) (*59*).

**Fig. 5.**
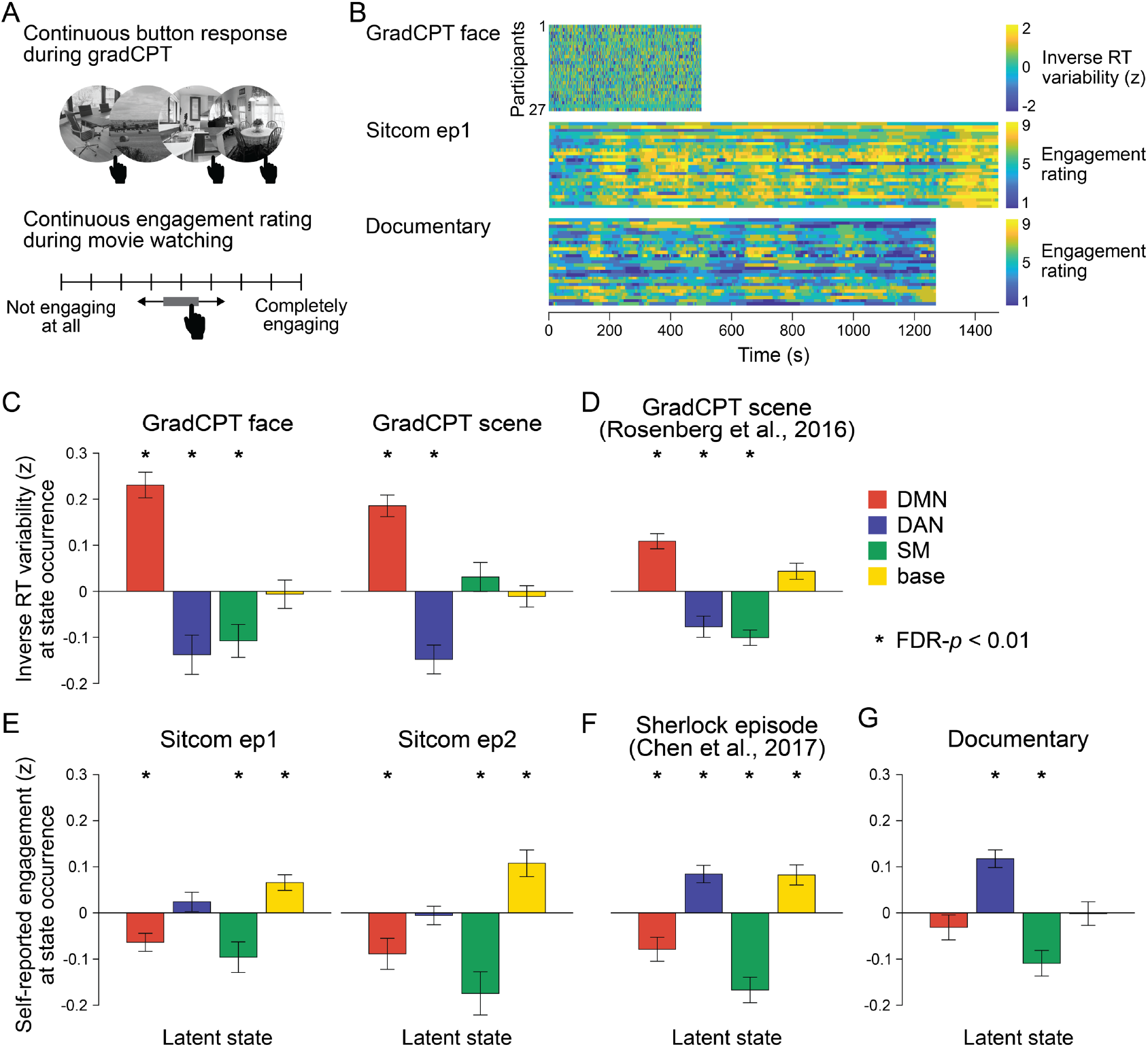
Relationship between latent neural states and attentional engagement. **(A)** Schematic illustration of the gradCPT and continuous narrative engagement rating. (Top) Participants were instructed to press a button at every second when a frequent-category image of a face or scene appeared (e.g., indoor scene), but to inhibit responding when an infrequent-category image appeared (e.g., outdoor scene). Stimuli gradually transitioned from one to the next. (Bottom) Participants re-watched the sitcom episodes and documentary after the fMRI scans. They were instructed to continuously adjust the scale bar to indicate their level of engagement as the audiovisual stimuli progressed. **(B)** Behavioral measures of attention in three fMRI conditions. Inverse RT variability was used as a measure of participants’ attention fluctuation during gradCPT. Continuous ratings of subjective engagement were used as measures of attention fluctuation during sitcom episodes and documentary watching. Both measures were *z*-normalized across time during the analysis. **(C-G)** Degrees of attentional engagement at moments of latent state occurrence. The attention measure at every time point was categorized into which latent state occurred at the corresponding moment and averaged per neural state. The bar graphs indicate the mean of these values across participants. The mean values were compared with the null distributions in which the latent state dynamics were circular-shifted (asterisks indicate FDR-*p* < 0.01). **(C, E, G)** Results of the fMRI runs in the SONG dataset. **(D, F)** The HMM trained on the SONG dataset was applied to decode the latent state dynamics of **(D)** the gradCPT data by Rosenberg et al. (*85*) (N=25), and **(F)** the Sherlock television watching data by Chen et al. (*64*) (N=16).

We asked whether neural state occurrence reflected participants’ attentional states. For each participant, we averaged time-resolved measures of attention based on the latent neural states that occurred at particular moments of time.

#### Distinct states correspond to engaged attention during tasks and movies

Different brain states accompanied successful task performance and engaged movie watching. During the gradCPT, participants were in a high attentional state when the DMN state occurred (**Fig. 5C**). Results replicated when the SONG-trained HMM was applied to the gradCPT data collected by Rosenberg et al. (*85*) (**Fig. 5D**). This finding conceptually replicates previous work that showed the DMN involvement during in-the-zone moments of the gradCPT (*26, 86*) and supports the role of the DMN in automated processing of both the extrinsic and intrinsic information (*83, 87, 88*).

Other neural states indicated moments of high attention during movie watching. During comedy sitcoms, the base state was associated with engaged attention (**Fig. 5E**). Results replicated when the SONG-trained HMM was applied to television episode watching data collected by Chen et al. (*64*) (N=16) (**Fig. 5F**). To our knowledge, the involvement of the base state at engaging moments of movie watching has not been reported previously. During the educational documentary, on the other hand, the DAN state was associated with engaged attention (**Fig. 5G**). When watching a less engaging but information-rich documentary, focusing may require goal-directed and voluntary control of attention (*34*). Together, the results imply that different neural states indicate engaged attention in different contexts.

#### A common state underlies attention lapses during tasks and movies

In contrast to moments of engaged attention, moments of attention lapses were associated with the same brain state during gradCPT performance and movie watching. The SM state occurred during moments of poor gradCPT performance in the SONG (**Fig. 5C**) and Rosenberg et al. (*85*) datasets (**Fig. 5D**). It also occurred during periods of disengaged focus on the comedy sitcoms (**Fig. 5E**), the television episode of Chen et al. (*64*) (N=16) (**Fig. 5F**), and the educational documentary (**Fig. 5G**). Higher head motion was observed during the SM state compared to the three other states (**fig. S11**). However, the latent states consistently predicted attention when head motion was included as a predictor in a linear model (main effect of HMM latent states, F > 3, *p* values < 0.05 for 7 fMRI runs in **Fig. 5C-G**; whereas the effect of head motion was inconsistent), demonstrating that the effects were not driven by motion alone.

To further investigate the role of the SM state, we applied the trained HMM to two external datasets, one containing gradCPT runs interleaved with fixation blocks (*85*), and the other containing working memory task runs interleaved with fixation blocks (*36, 37*). In both the gradCPT and working memory task, the SM state occurred more frequently during intermittent rest breaks in between the task blocks, whereas the DMN, DAN, and base states occurred prominently during the task blocks (**fig. S15**). These results suggest that the SM state indicates a state of inattention or disengagement common across task contexts.

## Discussion

Our study characterizes large-scale human fMRI activity as a traversal between latent states in a low-dimensional state space. Neural states spanned the center and extremes of predefined spatial gradient axes, with the state at the center functioning as a transitional hub. These gradients explained significant variance in neural dynamics, suggesting their role as a general latent manifold shared across cognitive processes. Global desynchronization marked moments of neural state transitions, with decreases in cofluctuation of the pairwise functional networks preceding state changes. The same latent states recurred across fMRI runs and independent datasets, with distinct state-traversal patterns during rest, task, and naturalistic conditions. Neural state dynamics were synchronized across participants during movie watching and temporally aligned to narrative event boundaries. Whereas *different* neural states were involved in attentionally engaged states in task and naturalistic contexts, a *common* neural state indicated inattention in both contexts. Together, our findings suggest that human cognition and attention arise from neural dynamics that traverse latent states in a shared low-dimensional gradient space.

Using a dynamical systems approach, cognitive neuroscientists have theorized that hierarchically modular systems of the brain communicate and process information dynamically (*3*). This framework, which characterizes the dynamics of systems-level interactions as a trajectory within a state space, has opened a new avenue to understanding the functional brain beyond what could be revealed from the univariate activity of local brain regions or their pairwise connections alone (*4*). Although dynamical systems approach has been adopted in non-human studies to understand behavior during targeted tasks (*89–92*), there is still a lack of understanding of how human cognition arises from the systems-level interactions, with a particularly sparse understanding on what gives rise to naturalistic, real-world cognition.

Our study adopted the assumption of low dimensionality of large-scale neural systems, which led us to intentionally identify states underlying whole-brain dynamics captured by the time series of only 25 parcels, which were then replicated with a higher number of parcels (**fig. S4**). We demonstrate that cognitive and attentional state fluctuations that occur over tens of seconds or minutes are represented globally in a low-dimensional state space. However, we acknowledge that depending on granularities of cognitive processes, the optimal dimensionality for representation and computation would differ as well as for the spatiotemporal scales at which the neural systems are involved.

Using fMRI data collected in rest, task, and naturalistic contexts, we identified four latent states that tile the principal gradient axes of functional brain connectome. Are these latent states—the DMN, DAN, SM and base states—generalizable “canonical” states of the human brain? When the HMM was applied to data from each condition separately, the inferred latent states differed (**fig. S2**). However, when the HMM was applied to datasets including diverse fMRI conditions like the SONG and HCP, the four states consistently reappeared, regardless of analytical choices (**figs. S1, S5**, and **S6**). Moreover, previous studies that applied different clustering or dimensionality reduction algorithms to fMRI data extracted qualitatively similar latent states or components (*14*). These studies and ours report latent axes that follow a long-standing theory of large-scale human functional systems (*93*). Neural dynamics span principal axes that dissociate unimodal to transmodal, and sensory to motor information processing systems. We propose a framework that can unify these observations and theories: the large-scale neural dynamics traverse canonical latent states in a low-dimensional subspace captured by the principal gradients of functional brain organization.

Previous studies reported functional relevance of latent state dynamics during controlled (*1, 20–22, 94*) and naturalistic tasks (*23, 24*). The current study aimed to unify these findings by generalizing the latent state model to multiple fMRI runs and datasets spanning rest, task, and naturalistic contexts. Intriguingly, the latent states commonly occurred in every scan type (**Fig. 3B**) but their functional roles differed depending on context. For example, during monotonous tasks that required constant exertion of sustained attention, the DMN state accompanied successful, stable performance whereas the DAN state characterized suboptimal performance (**Fig. 5C** and **D**). The antagonistic activity and functional relationship between the DMN and DAN has been reported in past studies that used resting-state (*31, 33*) or task fMRI (*26, 86, 95*). In contrast, in naturalistic contexts, the DMN state indicated low attentional engagement to narratives (**Fig. 5E** and **F**) and tended to follow event boundaries (**Fig. 4A** and **B**). The DAN state, on the other hand, indicated high attentional engagement at documentary watching condition (**Fig. 5G**) and was not modulated by event boundaries (**fig. S13A** and **B**). Our results indicate that the functional relationship between the DMN and DAN states shows more nuanced dependence to contexts.^1^ The findings highlight the need to probe both the controlled and naturalistic tasks with dense behavioral sampling to fully characterize the functional roles of these neural states (*96*).

In contrast to the context-specific DMN and DAN states, the SM state consistently indicated inattention or disengagement. The SM state occurred during poor task performance and low narrative engagement (**Fig. 5**) as well as during intermittent task breaks (**fig. S15**). The result implies that whereas the optimal neural state may vary with information processing demands, a suboptimal state is shared across contexts.

Finally, we propose that the base state, a transitional hub (**Fig. 2B**) at the center of the gradient subspace (**Fig. 1D**), acts as a state of natural equilibrium. Transitioning to the DMN, DAN, or SM states reflects incursion away from natural equilibrium (*2, 18*), as the brain enters functionally modular state. Intriguingly, the base state indicated high attentional engagement (**Fig. 5E** and **F**) and exhibited the highest occurrence proportion during naturalistic movie watching (**Fig. 3B**), whereas its functional involvement was comparatively minor during controlled tasks. This significant relevance to behavior verifies that the base state cannot simply be a byproduct of the model. We speculate that the base state may reflect a near-critical state that maximizes susceptibility to both the external and internal information—allowing for roughly equal weighting of both sides so that they can be integrated to form a coherent representation of the world—at the expense of stability of a certain functional network (*97–99*). When processing rich narratives, and especially when a person is fully immersed without having to exert cognitive effort, a less modular state that has high degrees of freedom to reach other states may be more likely to be involved.

This work provides a framework for understanding large-scale human brain dynamics and their relevance to cognition and behavior. Neural dynamics can be construed as traversals across latent states along the low-dimensional gradients, driven by interactions between functional networks. The traversals occur adaptively to external and internal demands, reflecting ongoing changes of cognition and attention in humans.

## Materials and Methods

### SitcOm, Nature documentary, Gradual-onset continuous performance task (SONG) neuroimaging dataset

#### Participants

Twenty-seven participants were recruited in Korea (all native Korean speakers; 2 left-handed, 15 females; age range 18–30 years with mean age 23 ± 3.16 years). Participants reported no history of visual, hearing, or any form of neurological impairment, passed the Ishihara 38 plates color vision deficiency (CVD) test^2^ for red-green color blindness, provided informed consent before taking part in the study, and were monetarily compensated. The study was approved by the Institutional Review Board of Sungkyunkwan University. None of the participants were excluded from analysis.

#### Study overview

Participants visited twice for a 3-hour experimental session per day. Sessions were separated by approximately one week on average (mean 8.59 ± 3.24 days, range 2 to 15 days). Two participants returned for an additional scan and behavioral session because technical difficulties prevented them from completing the experiment within the two days.

During the first scan session, participants watched the first episode of a sitcom as well as a documentary clip during fMRI. Scan order was counterbalanced. One participant’s sitcom episode 1 fMRI run was not analyzed because the data were not saved. Structural T1 images were collected after EPI acquisitions. Immediately after the MRI scan session, participants completed behavioral tasks in a different room. They first completed free recall of the two movie clips, in the order of viewing. These data are not analyzed here but were used to confirm that the participants were awake during the scans. Participants then were asked to complete continuous engagement ratings while re-watching the same audiovisual stimuli in the same order. This fMRI experiment lasted approximately 1 hour and the post-scan behavioral experiment lasted approximately 1.5 to 2 hours.

The second scan session began with two 10-min resting state runs in which the participants were asked to fixate on a centrally presented black cross on a gray background. Next, participants completed the gradCPT with face images, watched the second sitcom episode, and performed the gradCPT with scene images. After fMRI, participants completed two runs of a recognition memory task for the scene images that were viewed during gradCPT and a free recall of the sitcom episode 2. These data were not analyzed here. The continuous engagement rating for the second sitcom episode was not collected during this session due to time limitations. 18 of the 27 participants returned to the lab to complete the continuous engagement rating task for the second sitcom episode. The fMRI experiment lasted approximately 1.5 hours and the post-scan behavioral experiment lasted approximately 1 hour.

#### Sitcom episodes and documentary watching

##### Stimuli

Two episodes of the comedy sitcom and one educational documentary clip were used as audiovisual stimuli. The sitcom, *High kick through the roof*, is a South Korean comedy sitcom that was aired in 2009-2010 on a public television channel, MBC.^3^ The duration of the first episode was 24 m 36 s and the second episode was 24 m 15 s. These episodes were chosen because the narrative followed an interleaved ABAB sequence. Events of a story A (e.g., which took place in a forest and centered around two sisters) happened independently from events of a story B (e.g., which took place in a city and followed members of a large family), and the two storylines occurred in different times and places and included different characters. To avoid a transient increase in fMRI activity upon a sudden video presentation, we included a 30 s of a dummy video clip from the Minions Mini Movies (2019)^4^ prior to the presentations of the sitcom episodes which was discarded in analysis.

*Rivers of Korea, Part 1* (21 m 33 s) is a documentary that aired in 2020 by a public educational channel EBS Docuprime in South Korea. The documentary introduces the history and geography of the two largest Korean rivers, the Han and Nakdong Rivers. This stimulus was chosen because while it has rich and dynamically changing audiovisual and narrative content, it elicits an overall low degree of engagement due to its educational purpose (although individuals may vary in the degree to which they find it engaging). The first 22 s of the documentary was not included in the analysis to account for a sudden increase in brain activity upon video presentation.

The audiovisual stimuli were presented at a visual dimension of 1280 × 720 and frame rate of 29.97 Hz on a black background. 30 s of center fixation was included at the end of every naturalistic stimulus run. No additional task was given to participants during fMRI except for an instruction to stay vigilant and attentive to the video.

##### Continuous engagement rating

Participants re-watched the videos in a behavioral testing room while they were instructed to continuously adjust the scale bar from scale of 1 (Not at all engaging) to 9 (Completely engaging) which was visible on the bottom of the monitor. Participants were instructed to report their experience as closest to when they have watched the stimulus during the fMRI. The definition of engagement was given to participants following Song et al. (*59*) as: I find the story engaging when i) I am curious and excited to know what’s coming up next, ii) I am immersed in the story, iii) My attention is focused on the story, iv) The events are interesting; whereas I find the story not engaging when i) I am bored, ii) Other things pop into my mind, like my daily concerns or personal events, iii) My attention is wandering away from the story, iv) I can feel myself dozing off, v) The events are not interesting.

Participants were encouraged to adjust the scale bar whenever their subjective engagement changed during the sitcom episodes or documentary. All participants completed a practice session with a clip from a Korean Youtube channel.^5^ Stimuli were presented with Psychopy3 (*100*) on a MacBook 13-inch laptop. Participants were given freedom to turn on or turn off the light or have the headphone or speaker on. Upon completion of continuous engagement ratings, participants were asked to give an overall engagement score of the stimulus using the same 1 to 9 Likert scale.

Continuous engagement rating time courses, ranging from 1 to 9, was *z*-normalized across time per participant and convolved with the canonical hemodynamic response function to be related with the neural state dynamics.

#### Gradual-onset continuous performance task (gradCPT)

##### Task with face images

Grayscale face images (9 female and 1 male unique faces) were selected from the MIT Face Database (*27, 101*), cropped to a circle at a visual dimension of 300 × 300, and presented on a gray background. Five-hundred trials (450 female and 50 male face trials) were included in the run, with each unique face image appearing 50 times in a random sequence. No repeats of the images were allowed on consecutive trials. On each trial, an image gradually transitioned from one to the next using a linear pixel-by-pixel interpolation. The transition took 800 ms and the intact face image stayed for 200 ms when fully cohered. The task was to press a button on each trial when a female face appeared (90% of trials) but to inhibit making a response when a male face appeared (10% of trials). A fixation cross appeared for the first 1 s of the run, and the trial sequence started with a dummy stimulus (scrambled face). The run ended with a 10 s of center-fixation and lasted 8 min 33 s in total. A practice session was completed with the same face images prior to the scans.

##### Task with scene images

Colored scene images (300 indoor, 300 outdoor) were selected from the SUN database (*70, 102*). The stimulus was presented at a visual dimension of 500 × 500 on a gray background. Each individual saw 360 trial-unique images in a random order, with 300 (83.33%) coming from a frequent category (e.g., indoor) and 60 (16.67%) from an infrequent category (e.g., outdoor). Whether the indoor or outdoor scenes corresponded to the frequent category was counterbalanced across participants. Images transitioned in a pixelwise interpolation, with a transition occurring over 500 ms and the intact image lasting 700 ms. Participants were asked to press ‘1’ for frequent-category and ‘2’ for infrequent-category scene images. A fixation cross appeared at first 1 s of the run, and the trial started with a dummy stimulus (scrambled scene). The run ended with a 10 s of center-fixation and lasted 7 min 25 s in total. A practice session was completed with a different set of scene images prior to the scans.

##### Response time assignment algorithm

Each gradCPT trial included moments when an image was interpolated with the previous trial’s image followed by the fully cohered image. A maximum of two responses were recorded per trial. For most of trials, a single response or no response was recorded within the trial time window. However, if two responses were recorded in a trial (2.35% and 0.27% of all trials from tasks with face and scene images) and the response for the previous trial was missing, then the first response was regarded as a response for the previous trial and the second response as the response for the current trial. In cases when two responses were recorded but the response for the previous trial was not missing, or when the response for a single response trial happened before 40% of image coherence, we chose a response that favored a correct response.

##### Response time variability

To calculate an RT variability time course for each gradCPT run, the response times for incorrect and no-response trials were treated as NaNs, which were then filled by 1D linear interpolation. The RT time course was linearly detrended, and RT variability was calculated by taking the deviance from the mean RT at every TR. The RT variability time course was *z*-normalized across time within run and convolved with the canonical hemodynamic response function to be related with the neural state dynamics.

### FMRI image acquisition and preprocessing

Participants were scanned with a 3T scanner (Magnetom Prisma; Siemens Healthineers, Erlangen, Germany) with a 64-channel head coil. Anatomical images were acquired using a T1-weighted magnetization-prepared rapid gradient echo pulse sequence (repetition time [TR] = 2,200 ms, echo time [TE] = 2.44 ms, field of view = 256 mm × 256 mm, and 1 mm isotropic voxels). Functional images were acquired using a T2*-weighted echo planar imaging (EPI) sequence (TR = 1,000 ms, TE = 30 ms, multiband factor = 3, field of view = 240 mm × 240 mm, and 3 mm isotropic voxels, with 48 slices covering the whole brain). The number of TRs per run are as follows: resting-state run 1 (602 TR) and run 2 (602 TR), gradCPT with face (513 TR) and scene images (445 TR), sitcom episode 1 (1516 TR), episode 2 (1495 TR), and documentary (1303 TR). Visual stimuli were projected from a Propixx projector (VPixx Technologies, Bruno, Canada), with a resolution of 1920 × 1080 pixels and a refresh rate of 60 Hz. Auditory stimuli were delivered by MRI compatible in-ear headphones (MR Confon; Cambridge Research Systems, Rochester, UK).

Structural images were bias-field corrected and spatially normalized to the Montreal Neurological Institute (MNI) space using FSL. The first two images of the resting-state and gradCPT runs, 30 for sitcom episodes and 22 for documentary were discarded to allow the MR signal to achieve T1 equilibration. Functional images were motion-corrected using the six rigid-body transformation parameters. The functional images were intensity-normalized, and the FMRIB’s ICA-based X-noiseifier (FIX) was applied to automatically identify and remove noise components (*103–105*). The images were registered to MNI-aligned T1-weighted images. We additionally regressed out low-frequency components (high-pass filtering, *f* > 0.009 Hz), linear drift, and the global signal. The raw datasets of the SONG, Chang et al. (*66*), Rosenberg et al. (*85*), and Chen et al. (*64*) were all preprocessed with the same pipeline. Results replicated when processing included band-pass filtering in place of high-pass filtering (0.009 < *f* < 0.08 Hz) and when preprocessing did not include global signal regression (**fig. S5**). All analyses were conducted in volumetric space.

### Human Connectome Project (HCP) dataset

We used 3-Tesla and 7-Tesla data from 184 young adult participants in the HCP dataset. 3-Tesla data included 4 resting-state runs (*REST1* and *REST2* in the left-to-right and right-to-left phase encoding directions) and 2 runs each of the 7 tasks (*EMOTION, GAMBLING, LANGUAGE, MOTOR, RELATIONAL, SOCIAL*, and *WM*). 7-Tesla data included 4 resting-state runs (*REST1_PA, REST2_AP, REST3_PA, REST4_AP*) and 4 movie-watching runs (*MOVIE1_AP, MOVIE2_PA, MOVIE3_PA, MOVIE4_AP*). The combined data included 8400 TRs of the resting-state runs, 3880 TRs of task runs, and 3655 TRs of the movie-watching runs per participant. Of these 184 individuals, we excluded 19 who had not completed any of the scan runs, and 6 whose scan run was aborted earlier than others. Additionally, 40 participants’ data were discarded due to excessive head motion; having at least one fMRI run with more than 20% of the time points’ framewise displacement (FD) ≥ 0.5 or mean FD ≥ 0.5. This resulted in the analysis of 119 participants in total. We downloaded the MNI-aligned, minimally preprocessed structural and functional MRI images from the HCP repository (*106*). Additionally, the global signal, white matter and cerebrospinal fluid time courses, 12 head motion parameters (provided by Movement_Regressors.txt), and a low-frequency component (high-pass filtering, *f* > 0.009 Hz) were regressed from the data. For details on fMRI image acquisitions and task procedures, see (*36–38*).

### Hidden Markov model (HMM)

We used the HMM to characterize the dynamics of latent neural states that underlie large-scale functional brain activity (*28*). First, we parcellated the whole brain into 17 cortical networks (*29*) and 8 subcortical regions (*30*) and averaged the BOLD time series of the voxels that corresponded to these parcels. The 25 parcel time-courses of all participants’ every run were *z*-normalized within-run and concatenated.

Expectation-maximization (*107*) of the forward-backward algorithm was used to estimate the optimal model parameters: (i) the emission probability *p*(*y*_*t*_|*x*_*t*_) of the observed fMRI time series {*y*_1_…*y*_*T*_} from the hidden latent sequence {*x*_1_…*x*_*T*_}, and (ii) the first-order Markovian transition probabilities *p*(*x*_*t*_ = *s*_*i*_ |*x*_*t*−1_ = *s*_*j*_) for 1 ≤ *i, j* ≤ *K*. The emission probability was modeled using a mixture Gaussian density function (hmmlearn.hmm.GaussianHMM), such that 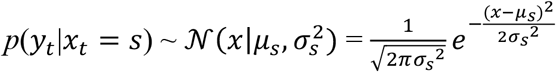 per discrete *K* number of latent state *s* ∈ {*s*_1_…*s*_*K*_}, where 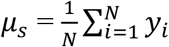 and 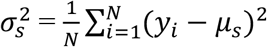 for a set of observed fMRI time steps {*y*_1_…*y*_*N*_} identified as the latent state *s*. The *µ*_*s*_ and *σ*_*s*_^2^ represent the mean activation and covariance patterns of the 25 parcels per each state (**Fig. 1A**). The transitions between the hidden states were assumed to have a form of a first-order Markov chain, such that if *a*_*ij*_ represents the probability of transitioning from state *i* to state *j*, then 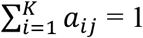. The inference procedure terminated if there was no longer a gain in log-likelihood during the re-estimation process of the forward-backward algorithm or if the number of maximum 1,000 iterations was reached. We initialized the HMM parameters using the output of k-means clustering to overcome the problem of falling into a local minima (sklearn.cluster.KMeans).

The estimated transition and emission probabilities were applied to decode the most probable latent state sequence conditioned on the observed fMRI time series using a Viterbi algorithm (*108*). The Viterbi algorithm estimates the probability of each latent state being the most likely state at a specific time point. We chose the state with the highest probability at every time step (TR), thus discretizing the latent sequence.

To choose the optimal number of latent states (K), a hyperparameter that needed to be selected in advance, we conducted the HMM in a leave-one-subject-out cross-validated manner where we trained the HMM on all participants but one to infer transition and emission probabilities, and applied the HMM to decode the latent state dynamics of the held-out participant. A Calinski-Harabasz score was compared across the choice of K from 2 to 10 (*19, 24, 109*) (**fig. S1**). The K with the largest mean Calinski-Harabasz score across cross-validations was selected, and we conducted the HMM on all participants’ data with the chosen K. The HMM inference and decoding procedure was repeated 10 times and the instance with the maximum expected likelihood was chosen as a final result.

The surrogate latent sequence was generated by having 25-parcel time series circular-shifted across time respectively for each parcel, thereby disrupting meaningful covariance between parcels while retaining temporal characteristics of the time series, and applying the same HMM fitting and decoding algorithms 1,000 times. The maximum number of estimations was set as the number of iterations that was reached during the actual HMM procedure (1,000 for SONG and 248 for HCP).

Unless otherwise noted, the HMM parameter inference was conducted on the SONG and HCP datasets respectively and the model decoded latent state sequence of the same dataset. However, for analyses validating the functional roles of the latent states, the parameters inferred from SONG data were used to decode the latent state sequence of external datasets collected by Chang et al. (*66*) (**Fig. 4B**), Rosenberg et al. (*85*) (**Fig. 5D, fig. S15A-B**), Chen et al. (*64*) (**Fig. 5F**), and WM runs of the HCP dataset (*36, 37*) (**fig. S15C-D**).

**Figure 1B** shows mean activity and covariance patterns derived from the Gaussian emission probability estimation. The brain surfaces were visualized with nilearn.plotting.plot_surf_stat_map. The parcel boundaries in **Fig. 1B** are smoothed from volume-to-surface reconstruction.

Covariance strength was operationalized as the sum of the absolute covariance weights of all possible pairwise edges. Covariance strength calculated for each latent state was compared to chance distributions generated from covariance matrices estimated from the HMMs conducted on the circular-shifted 25 parcel time series (1,000 iterations, two-tailed non-parametric permutation tests, FDR-corrected for the number of latent states).

### Functional gradients and principal component analysis

The gradients of the cortical and subcortical voxels estimated by Margulies et al. (*25*) were downloaded from the repository.^6^ The gradient values of the voxels within each 25 parcel were averaged to represent each parcel’s position in the gradient space. To situate the latent states in the Margulies et al.’s (*25*) gradient axes, we took the mean of elementwise-product between these gradient values and the mean activity loadings of the 25 parcels inferred by the HMM.

We evaluated how well these pre-defined gradients capture large-scale neural dynamics in the SONG and HCP datasets compared to data-driven principal components (PCs) defined in each dataset, respectively. To do so, we first conducted principal component analysis on the concatenated *z*-normalized parcel time series of all participants’ fMRI runs (sklearn.decomposition.PCA). The PC coefficients of the respective datasets were correlated with the mean gradient values of 25 parcels using Pearson’s correlations. The *r* value was compared to a null distribution in which gradient values were correlated with the null PC coefficients that were conducted on the circular-shifted time series (1,000 times, two-tailed non-parametric permutation tests, FDR-corrected for top-2 components that were tested).

Next, we computed the explained variance of the PCs and gradients as the mean of squared Pearson’s correlations (*r*^2^) between the 25 parcel time courses and the PC scores or gradient-projected time courses. The explained variance (*r*^2^) of the PCs and gradients, of all participants’ every run, were compared using paired *t*-tests, FDR-corrected for 2 comparisons. The mean *r*^2^ across all participants’ fMRI runs was compared to a null distribution generated by correlating circular-shifted 25 parcel time courses with the PC and gradient time courses (1,000 iterations, two-tailed non-parametric permutation tests, FDR-corrected for 2 comparisons).

### Cofluctuation time course time-aligned to neural state transitions

Cofluctuation, the absolute element-wise product of the normalized activity in two regions, was computed based on ref. (*45, 46*). We normalized the time courses of the two regions among the 25 parcels, *x*_*i*_ and *x*_*j*_ (*i* = 1…T and *j* = 1…T), such that 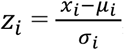, where 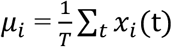 and 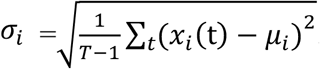. The cofluctuation time series between *z*_*i*_ and *z*_*j*_ was computed as the absolute of the element-wise product, |*z*_*i*_ · *z*_*j*_|, which represents the magnitude of moment-to-moment cofluctuations between region *i* and *j* based on their baseline activities. Cofluctuation was computed for every pair of parcels, resulting in a time-resolved, 25 (parcel) × 25 × T (number of TRs) matrix for each fMRI run, with a symmetric matrix at every time point.

We categorized the 25-parcel pairs to cortico-cortical (136 pairs), cortico-subcortical (136 pairs), and subcortico-subcortical (28 pairs) connection categories. The cofluctuation time courses were time-aligned to multiple moments of state transitions indicated by the HMM inference, which were averaged within a participant. The time-aligned mean cofluctuation of each pair was then averaged across all runs of the entire participants.

The chance distributions were created in two ways. First, cofluctuation time courses were time-aligned to the circular-shifted indices of neural state transitions (1,000 iterations; **Fig. 2A**). Second, we circular-shifted the 25-parcel time series, thereby disrupting their covariance structure, and conducted HMM inference on these null time series (1,000 iterations; **fig. S9**). The time-aligned cofluctuation of every pair of parcels was averaged per category and was compared to chance distributions using *z*-statistics and two-tailed non-parametric permutation tests. The significance at each time point was FDR-corrected for the number of time points (i.e., -3 to 3 from the onset of new latent states). Furthermore, to compare the degrees of cofluctuation change between cortico-cortical, cortico-subcortical, and subcortico-subcortical connection categories, mean cofluctuation difference at time *t-1* and *t+3* was taken per category and the values were compared across categories using paired Wilcoxon signed rank tests (FDR-corrected for three pairwise comparisons).

### Neural state transition probabilities

The *T-1* number of transitions in each participant’s latent state sequence were categorized based on which state it transitioned from (at *t-1*) and which state it transitioned to (at *t*) as a 4 (‘from’ state) × 4 (‘to’ state) transition-count matrix. We controlled for the number of state occurrences by either dividing each element by the sum of each row, which identified the probabilities of transitioning ‘to’ one of the four latent states (**Fig. 2B**), or dividing by the sum of each column, which identified the probabilities of transitioning ‘from’ one of the four latent states (**fig. S10**). The transition probability matrices estimated from every participant’s every run were averaged. The chance distribution was created by conducting the HMM fits on the circular-shifted 25 parcel time series (1,000 iterations). Significance was computed for each state pair of the transition matrix, using the two-tailed non-parametric permutation tests (FDR-corrected for 16 pairs).

### Global cofluctuation of the latent states

Cofluctuation time courses of all pairs of parcels were averaged within a run of each participant. This global cofluctuation measure at every TR was categorized based on the HMM latent state identification, which was then averaged per state (**Fig. 2C**). Cofluctuation values of the base state of all participants’ entire runs were compared with the DMN, DAN, and SM states’ using the paired *t*-tests (FDR-corrected for 3 comparisons). The values at each latent state were averaged and compared to a chance distribution in which the analysis was repeated with a circular-shifted latent state sequence (1,000 iterations, two-tailed non-parametric permutation tests, FDR-corrected for the number of states).

### Fractional occupancy of the latent states

Using the HMM-derived latent state sequence, we calculated fractional occupancy of the latent states for all participants’ every run. Fractional occupancy is the probability of latent state occurrence over the entire fMRI scan sequence, with a chance value of 25% when K = 4. The mean of all participants’ fractional occupancy values was bootstrapped (10,000 iterations) for visualization in **Fig. 3B**.

### Pairwise participant similarity of the latent state sequence

The similarity between pairs of participants’ latent state sequences was computed as the ratio of the times when the same state occurred over the entire time course. The mean similarity was compared to the chance distribution in which participants’ neural state dynamics were circular-shifted 1,000 times (FDR-corrected for the number of fMRI runs). To compare the degree of synchrony across conditions, we bootstrapped the same pairs of participants with replacement (*number of participants*C*2* iterations) in paired conditions and took the differences of bootstrapped participant pairs’ latent state sequence similarities. The median of these differences was extracted 1,000 times and the distribution was compared to 0 non-parametrically.

### Neural state dynamics at narrative event boundaries

Narrative event boundaries were marked by the experimenter at moments in the sitcom episodes when an event of a storyline transitioned to another event of a different story. Both sitcom episodes comprised 13 events (7 events of story A and 6 events of story B), and thus 12 event boundaries. The latent state sequences at *t-2* and *t+20* TRs from each of the 12 event boundaries were extracted and the mean probability of state occurrence across these event boundaries was computed for every latent state within a participant. The probabilities were then averaged across participants. The state occurrence probability at every time step was compared to a chance distribution that was created by relating neural state dynamics to circular-shifted moments of event boundaries (1,000 times, two-tailed non-parametric permutation tests, FDR-corrected for number of time points). An audio-story listening data of Chang et al. (*66*) comprised 45 interleaved events. Because TR resolution (TR = 1.5 s) was different from the SONG dataset, latent neural states from *t-2* to *t+16* TRs from event boundaries were used in analysis.

Next, we compared transitions made to the DMN state at event boundaries to transitions made to the DMN state at moments other than event boundaries. We categorized every transition to the DMN state (i.e., from the DAN, SM, or base state) based on whether it occurred 5 to 15 TRs (for Chang et al. (*66*), 4 to 12 TRs) after a narrative event boundary or not. The proportions of DAN-to-DMN, SM-to-DMN, and base-to-DMN state transitions at event boundaries and non-event boundaries were compared using paired Wilcoxon signed rank tests (FDR-corrected for three comparisons). The interaction between the DMN-preceding latent states and event boundary conditions was tested using the repeated measures ANOVA.

### Neural state dynamics related to attention dynamics

We measured participants’ attention fluctuations during gradCPT and movie-watching fMRI scans. Fluctuations during gradCPT performance were inferred from the inverted RT variability time course (*z*-normalized) collected concurrently during the gradCPT scans with face and scene images, as well as during gradCPT in the Rosenberg et al. (*85*) dataset. Continuous engagement rating time courses (*z*-normalized) collected after the sitcom episode and documentary watching scans were used to infer changes in the degree to which the naturalistic stimuli were engaging over time. Engagement ratings of the Sherlock dataset (*64*) were collected by Song et al. (*59*) from an independent group of participants.

To relate attentional dynamics to the occurrence of neural states, we categorized each person’s attention measure at every TR based on the HMM latent state identification, which was averaged per state. The mean attention measures of the four states were averaged across participants and compared to a chance distribution in which the attention measures were circular-shifted to be related to the latent state dynamics (1,000 iterations, two-tailed non-parametric permutation tests, FDR-corrected for the number of states). For Sherlock dataset only, a single group-average engagement time course was related to the latent state sequence of each fMRI participant because we did not have fMRI participant-specific behavioral ratings.

Linear mixed-effects models were conducted on the 7 fMRI runs (**Fig. 5C-G**), where the model predicted attention measure at every time step from the inferred HMM state indices and head motion (framewise displacement computed after fMRI preprocessing). The participant index was treated as a random effect. The significance of the two main effects and their interaction were computed using ANOVA.

To investigate the role of the SM state, we analyzed the gradCPT data collected by Rosenberg et al. (*85*) and two sessions of the HCP WM task (**fig. S15**). The Rosenberg et al. (*85*) dataset (N=25) included gradCPT task blocks separated by intervening fixation blocks. The WM task run of the HCP dataset included 2-back and 0-back WM task blocks and fixation blocks (N=119). Nine participants’ LR runs and 11 participants’ RL runs in the HCP WM dataset were discarded in the analysis either because their block orders differed from the other HCP participants’ or because the experiment log was not saved in the dataset repository. Latent state fractional occupancy values were computed for each task block within a participant. Comparisons of a state’s fractional occupancies across block types were based on paired *t*-tests (FDR-corrected for the number of states). The interaction between latent neural states and task block types was tested using repeated measures ANOVA.

## Acknowledgments

We thank Kyung Soo Song for suggestions on audiovisual narrative stimuli. We thank JeongJun Park for help with fMRI and behavioral data collection, Jeongwon Shin for help with behavioral data collection, Boohee Choi for help with fMRI data collection, and Wooyoung Cho and Sunghyun Ban for their help with recall transcriptions. We thank Kyeong-Jin Tark and members of the Awh & Vogel lab for sharing experiment code, and Jiwoong Park for sharing a movie annotation program. We thank Yuan Chang Leong, Seok-Jun Hong, Emily S. Finn, and JeongJun Park for helpful discussions and comments on the manuscript. We thank many researchers who open-sourced their data and code that was used in the project. Research was supported by resources provided by the University of Chicago Research Computing Center.

## Author contributions

Conceptualization: HS, WMS, MDR

Methodology: HS, WMS, MDR

Investigation: HS

Visualization: HS

Funding acquisition: WMS, MDR

Project administration: HS, WMS, MDR

Supervision: WMS, MDR

Writing – original draft: HS, MDR

Writing – review & editing: HS, WMS, MDR

## Data and materials availability

De-identified MRI data will be made available on OpenNeuro and behavioral data and experimental code will be shared on https://github.com/hyssong/neuraldynamics upon publication.

## Supplementary Materials

Figs. S1 to S15

Supplementary Text

## Supplementary Materials

**Fig. S1.**
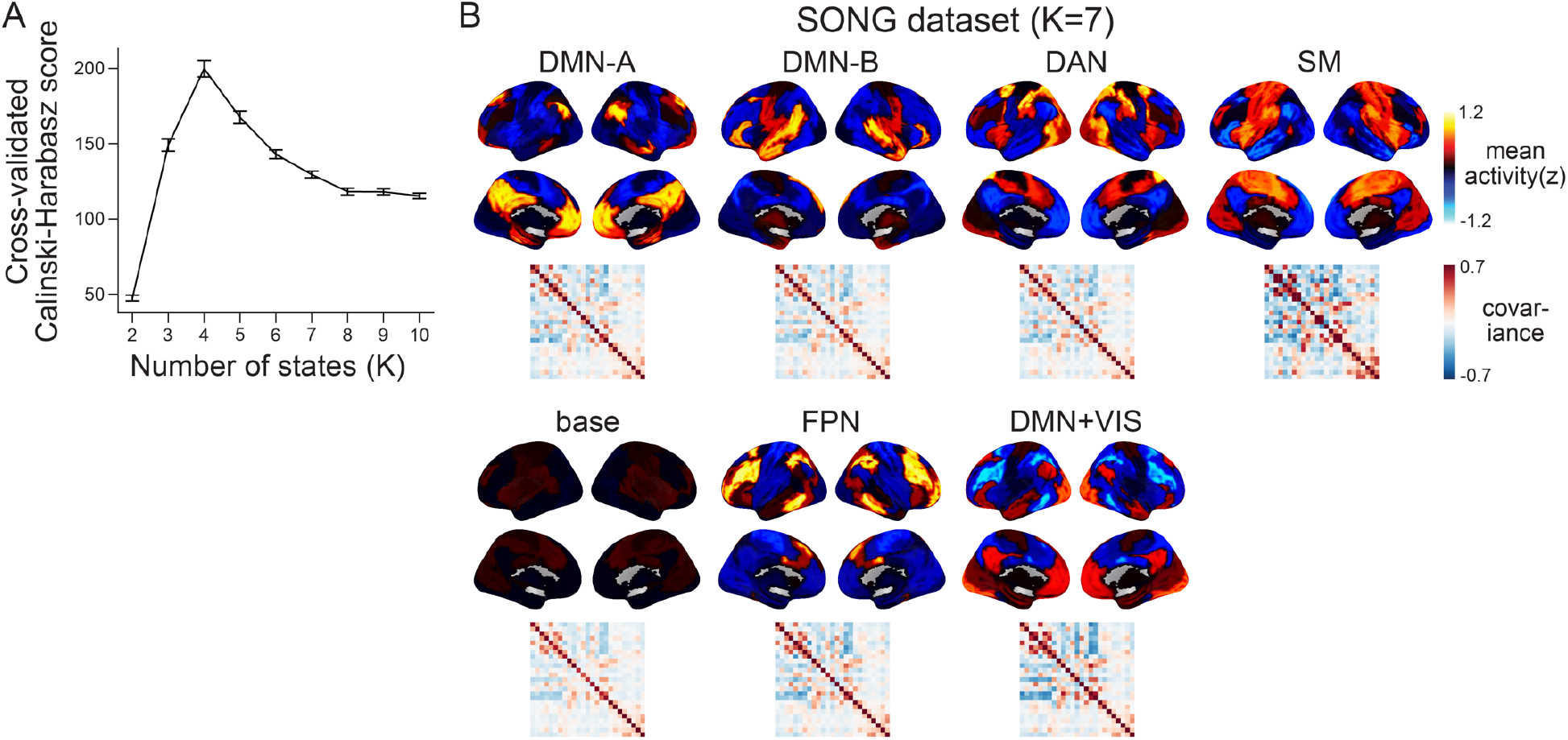
The choice of the number of states (K) in latent state inference. **(A)** To determine a value of K that optimizes HMM state inference from the SONG dataset, we iteratively calculated the Calinski-Harabasz scores using leave-one-subject-out cross-validation with K ranging from 2 to 10. Specifically, the HMM was trained on data from all participants but one to estimate parameters, emission and transition probabilities. The model was then applied to decode the latent state sequence of the held-out participant. With the held-out participant’s fMRI time series, the BOLD pattern similarity of the within-versus across-latent states were compared using the Calinski-Harabasz score, such that higher score indicates higher within-state cluster cohesion compared to the across-state dispersion (*19, 24, 109*). The line indicates the mean of the cross-validated Calinski-Harabasz scores and the error bars indicate S.E.M. **(B)** HMM latent neural states inferred from the SONG dataset using a different choice of K (K = 7). When more states were inferred, we observed subdivisions of and additions to the four neural states in **Fig. 1B**. The DAN, SM, and base states appeared with similar activity patterns (*r* = 0.678, 0.923, 0.445 respectively). The DMN state was subdivided into two DMN states. The DMN-A corresponded to the DMN core and the medial temporal subsystem. The DMN-B corresponded to the dorsal medial subsystem (*110*). Two additional neural states were inferred: one which exhibited the highest activity at the frontoparietal network (FPN) and the other showing activity in both the DMN and visual network (VIS).

**Fig. S2.**
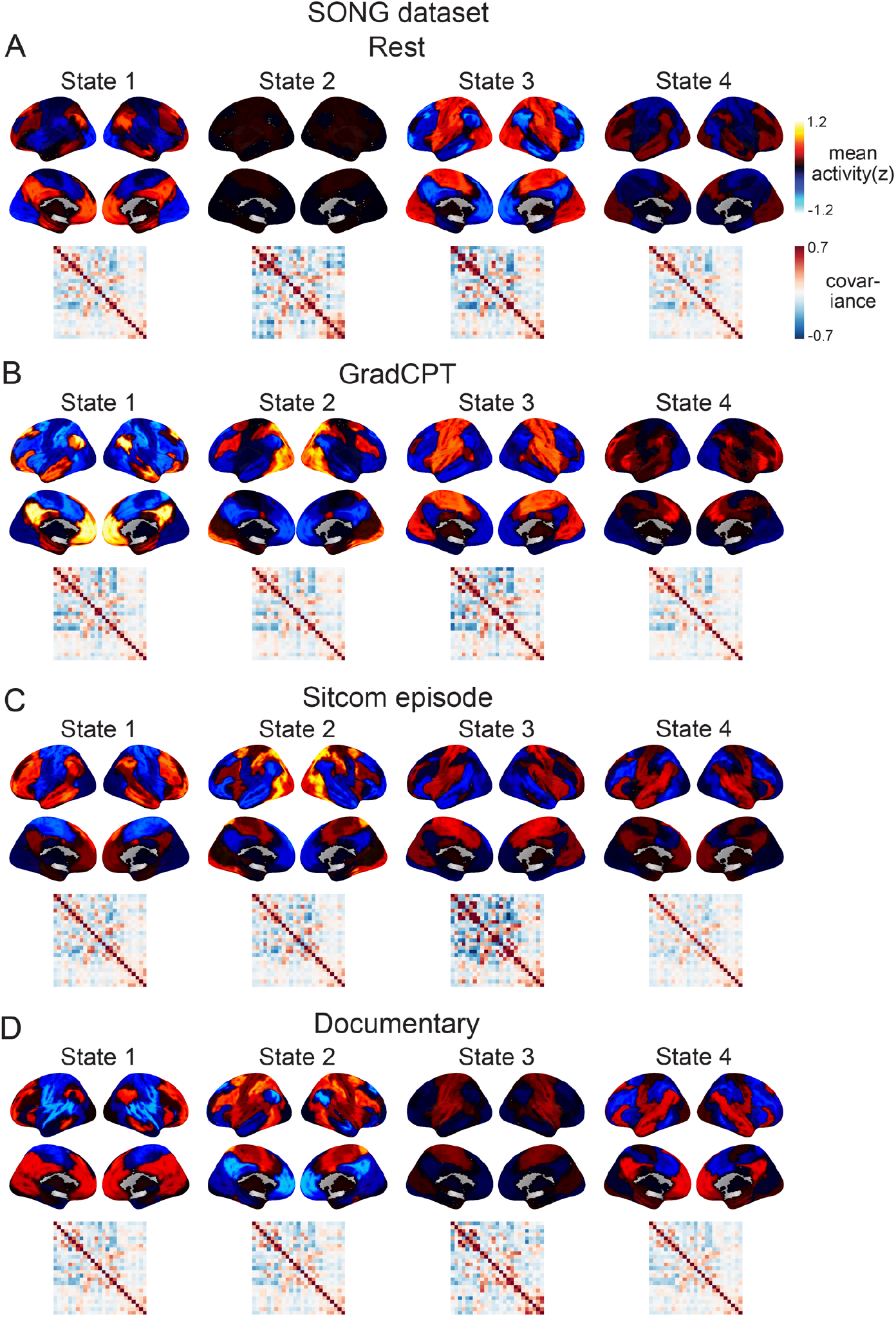
Latent state inference conducted separately to each condition of the SONG dataset. **(A)** *Rest* includes the two resting-state runs (600 TR × 2 for every 27 participant), **(B)** *GradCPT* includes the two gradCPT runs with face (511 TR) and scene (443 TR) images, **(C)** *Sitcom episode* includes the two sitcom-episode-watching runs (episode 1: 1486 TR, episode 2: 1465 TR), and **(D)** *Documentary* corresponds to the single documentary-watching run (1281 TR). The states are ordered based on the activation pattern similarity to the DMN, DAN, SM, and base states inferred from the full dataset (**Fig. 1B**). The pattern similarities are as follows in the order of DMN, DAN, SM, and base states: *Rest*: *r* = 0.801, 0.332, 0.722, 0.369; *GradCPT*: *r* = 0.956, 0.747, 0.928, 0.276; *Sitcom ep*: *r* = 0.486, 0.600, 0.737, 0.882; *Documentary*: *r* = 0.526, 0.847, 0.747, 0.663.

**Fig. S3.**
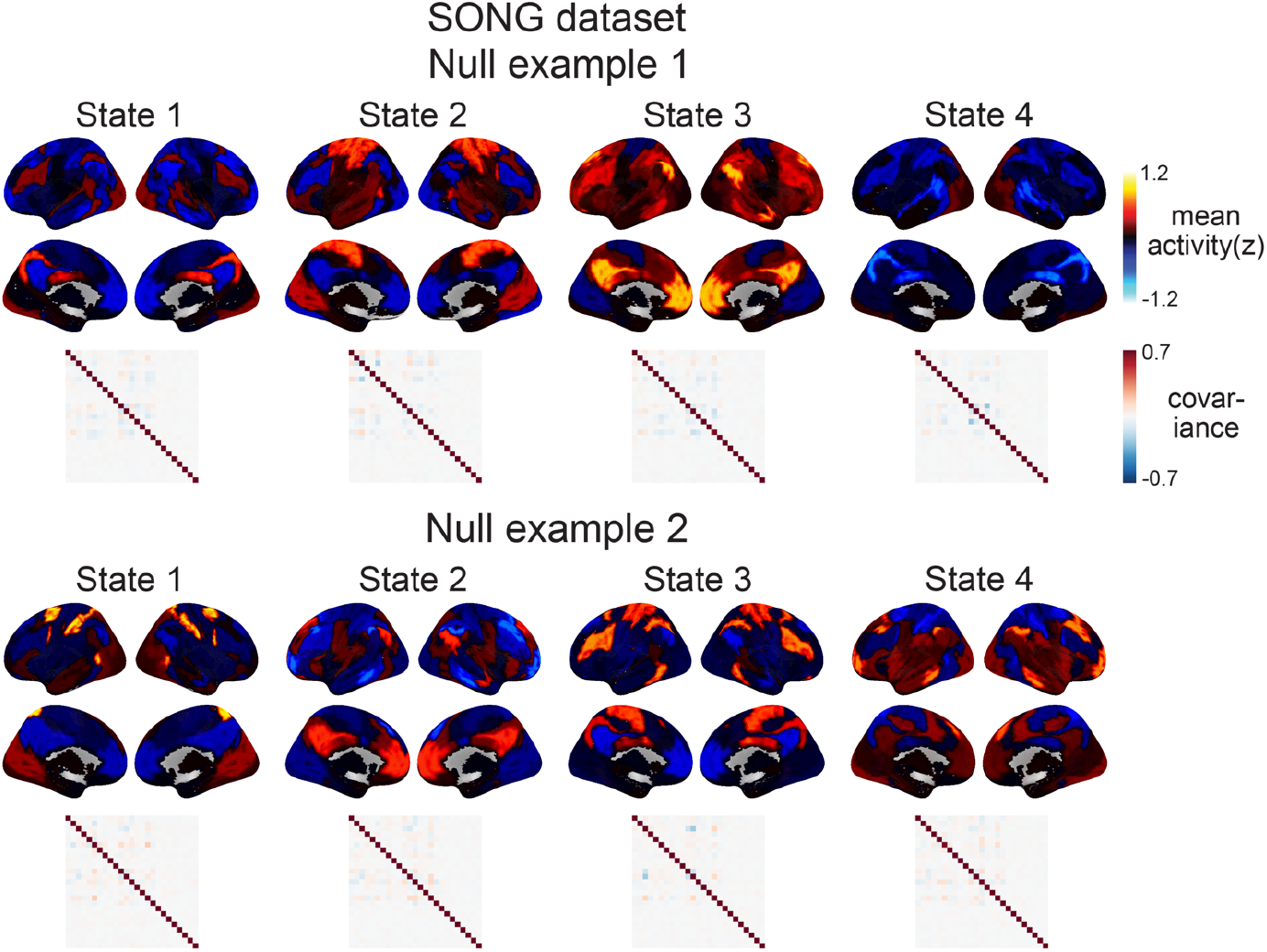
Examples of the null latent states derived from the HMMs conducted on the surrogate fMRI time series of the SONG dataset (1,000 iterations). The 25 parcel time courses from each fMRI run and participant were circular-shifted, separately for each parcel, which disrupts the covariance across parcels while retaining the temporal characteristics of each individual parcel. Compared to the latent states derived from the actual fMRI time series (**Fig. 1B**), these null states did not show systematic activity patterns and exhibited weaker covariance strengths. In **Fig. 1B**, the SM state exhibited the highest covariance strength (SONG: 52.570, HCP: 58.196), followed by comparable DMN (SONG: 37.277, HCP: 28.353) and DAN (SONG: 37.303, HCP: 29.525), whereas the base state exhibited the lowest covariance strength (SONG: 27.603, HCP: 25.426). On the contrary, the mean covariance strengths of the four surrogate covariance matrices were in the range of 3.215-4.432 for SONG and 0.986-2.204 for HCP datasets over 1,000 iterations.

**Fig S4.**
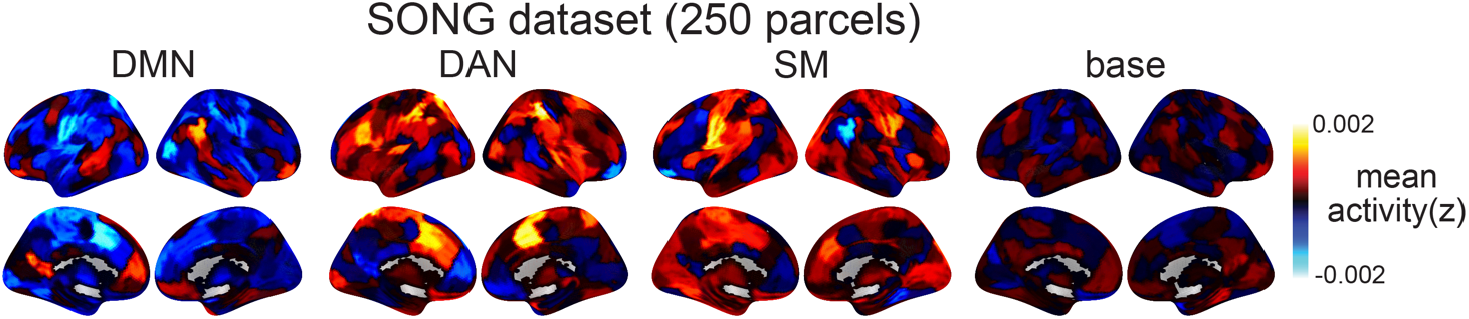
Latent state inference using a different whole-brain parcellation scheme. In the main text, we chose a fairly low-dimensional parcellation scheme (17 cortical and 8 subcortical) because the HMM produces a poorer fit when the dimensionality of the time series increases. The reason is that the number of parameters that need to be inferred for emission probability, *d*^2^ (*covariance*) + *d* (*mean*), increase exponentially with the increase in the number of parcel dimension *d*. Here, we test the robustness of the four latent states by using 200 cortical and 50 subcortical parcels, but applied dimensionality reduction to these 250 parcel time series prior to using them as inputs to the HMM. The matrices were randomly projected to 25-dimensional time series matrices by multiplying the time-by-250 matrix with the randomly generated 250-by-25 matrices. Random projection is a validated dimensionality reduction tool (*111*) that does not impose any constraints on how the latent dimensions should be (e.g., the principal component analysis should find latent dimensions that maximize explained variance and are orthogonal to one another). The HMM fit was conducted 500 times. For each iteration, after the HMM fit, we inverse-projected 25-dimensional mean activation patterns to 250 dimensions. The decoded latent state sequence of each iteration was compared to the decoded state sequence of the 25-parcel time series that we report in the manuscript. By finding the latent state sequence that shows the highest sequence similarity (mean 40.7 ± 3.45%, given a chance of 25%), we re-ordered the estimated mean activations and averaged across 500 iterations. The Pearson’s correlations between these average activation patterns and the inferred latent states in **Fig. 1B** are DMN: 0.549, DAN: 0.406, SM: 0.487, and base: 0.178. Thus, we see similar states in the SONG dataset even when input time series do not come from canonical functional networks. The DMN, DAN, SM, and base states are not specific to the parcellation scheme presented in the main text.

**Fig. S5.**
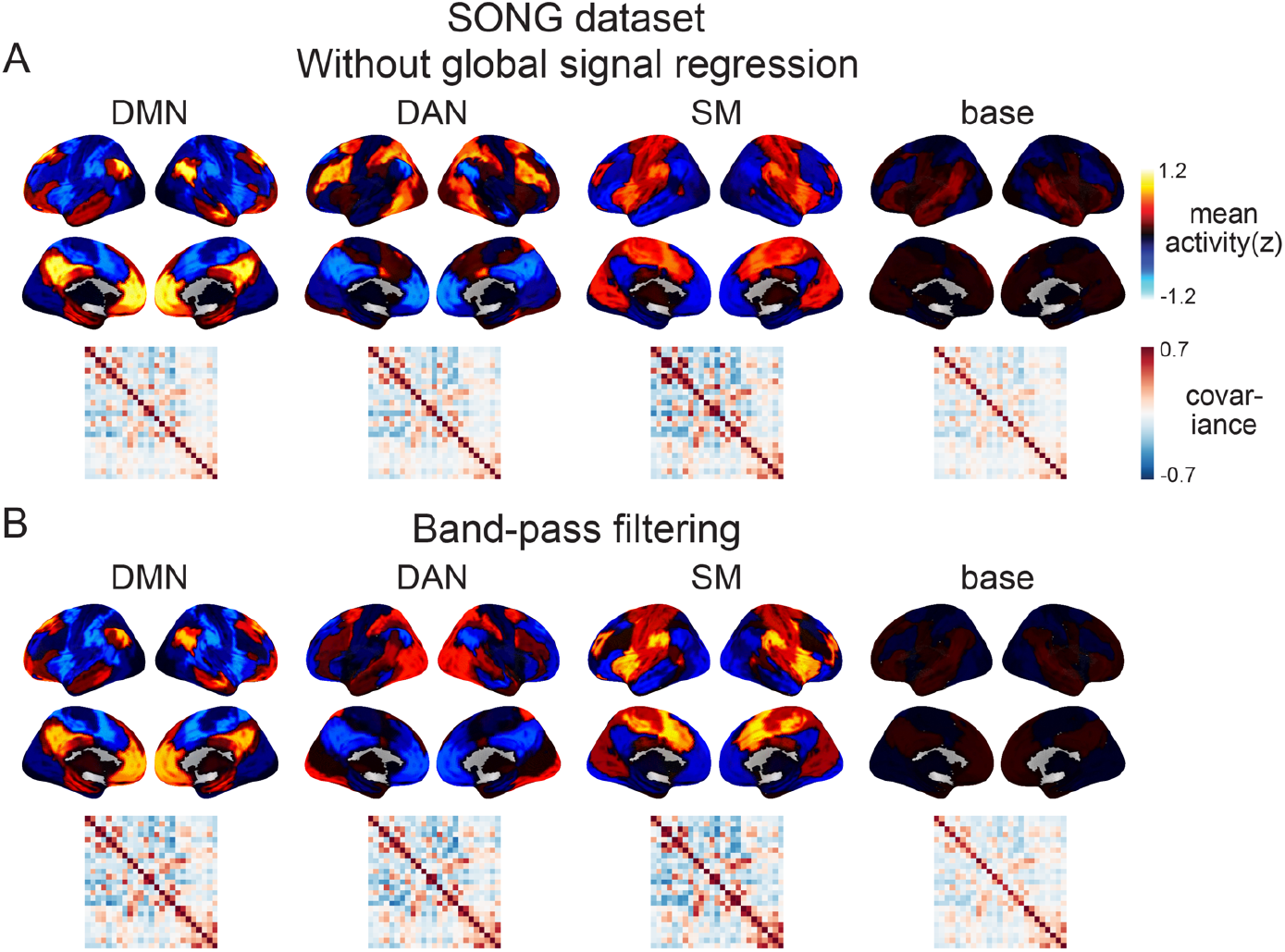
The inferred latent states and their dynamics are robust to the choice of the fMRI preprocessing approach. **(A)** We conducted HMM on preprocessed fMRI time series that did not undergo global signal regression. The output time series were highly similar with or without global signal regression, resulting in similar neural state dynamics (93.96% of the time points the same when assigned to the best-matching state, given a chance of 25%) and similar activity patterns of the four latent states (*r* = 0.999, 1.0, 0.997, 0.997, respectively). **(B)** We conducted HMM on the preprocessed fMRI time series that were temporally band-pass filtered (0.009 < *f* < 0.08 Hz) rather than high-pass filtered (*f* > 0.009 Hz). The inferred neural states had similar activity patterns (*r* = 0.961, 0.709, 0.883, 0.757), though their dynamics were not as similar as in **A** (28.59% of the time points the same). However, for all results reported in the article, we validated that the results were replicated in the band-pass filtered version of the analysis.

**Fig. S6.**
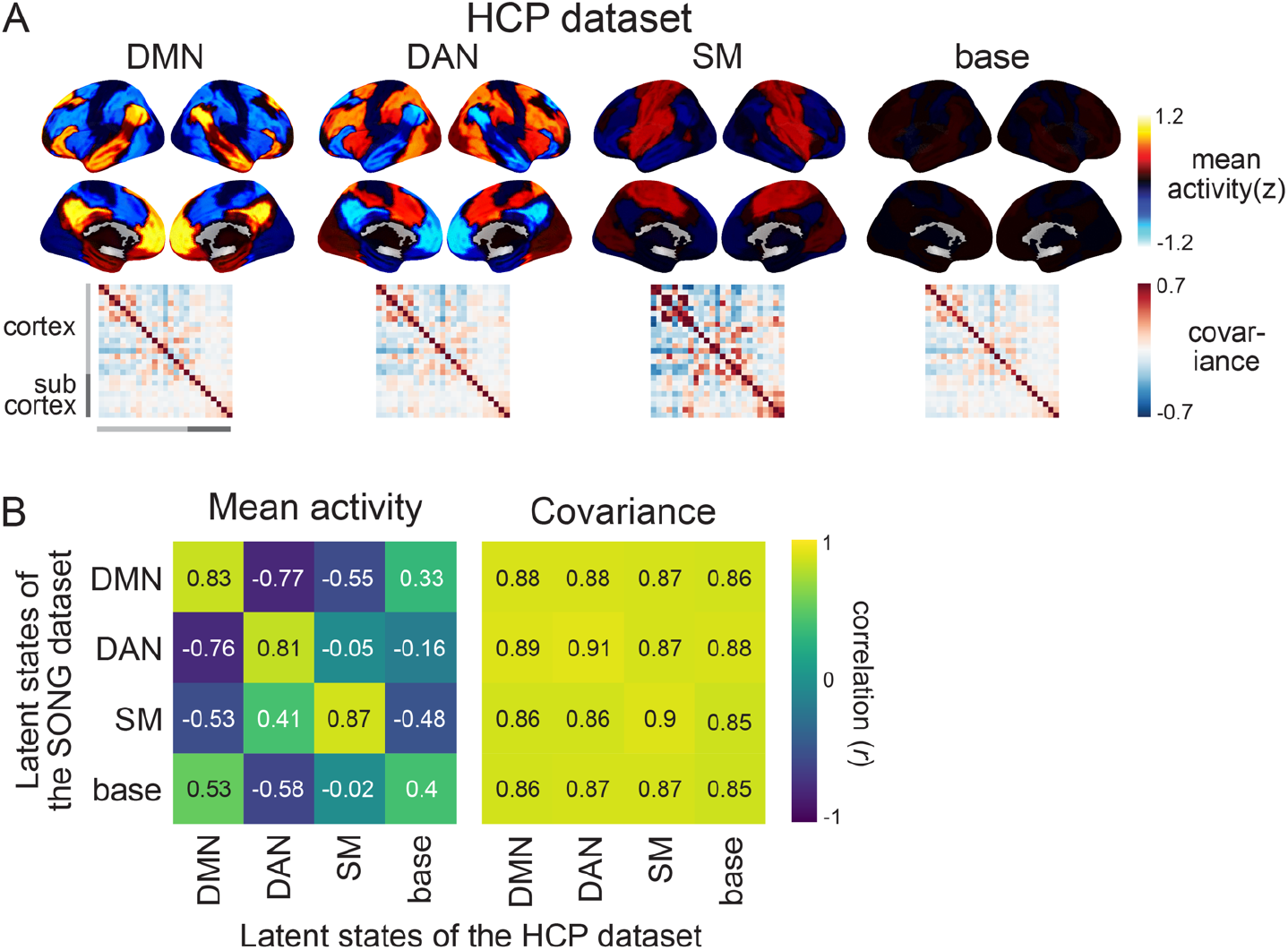
Latent state inference on the HCP dataset. **(A)** Four latent states inferred by the HMM fits to the HCP dataset. The figure complements **Fig. 1B. (B)** Comparison of the latent states inferred from the SONG (**Fig. 1B**) and HCP datasets. The DMN, DAN, and SM states showed similar mean activity patterns. We refrained from making interpretations about the base state’s activity patterns because the mean activity of most of the parcels was close to *z* = 0. The covariance patterns were similar throughout all pairwise neural states, although covariance strength values differed across neural states. To further validate the similarity of the SONG- and HCP-identified latent states, we trained the HMM on each dataset and applied it to decode the latent state sequence of the other dataset. When the SONG-trained HMM decoded the latent state sequence of the HCP dataset, 61.18% of the time points were the same as the latent state sequence identified from the HCP-trained HMM (with a chance of 25%). When the HCP-trained HMM decoded the latent state sequence of the SONG dataset, 61.67% of the time points were the same as the latent state sequence identified from the SONG-trained HMM.

**Fig. S7.**
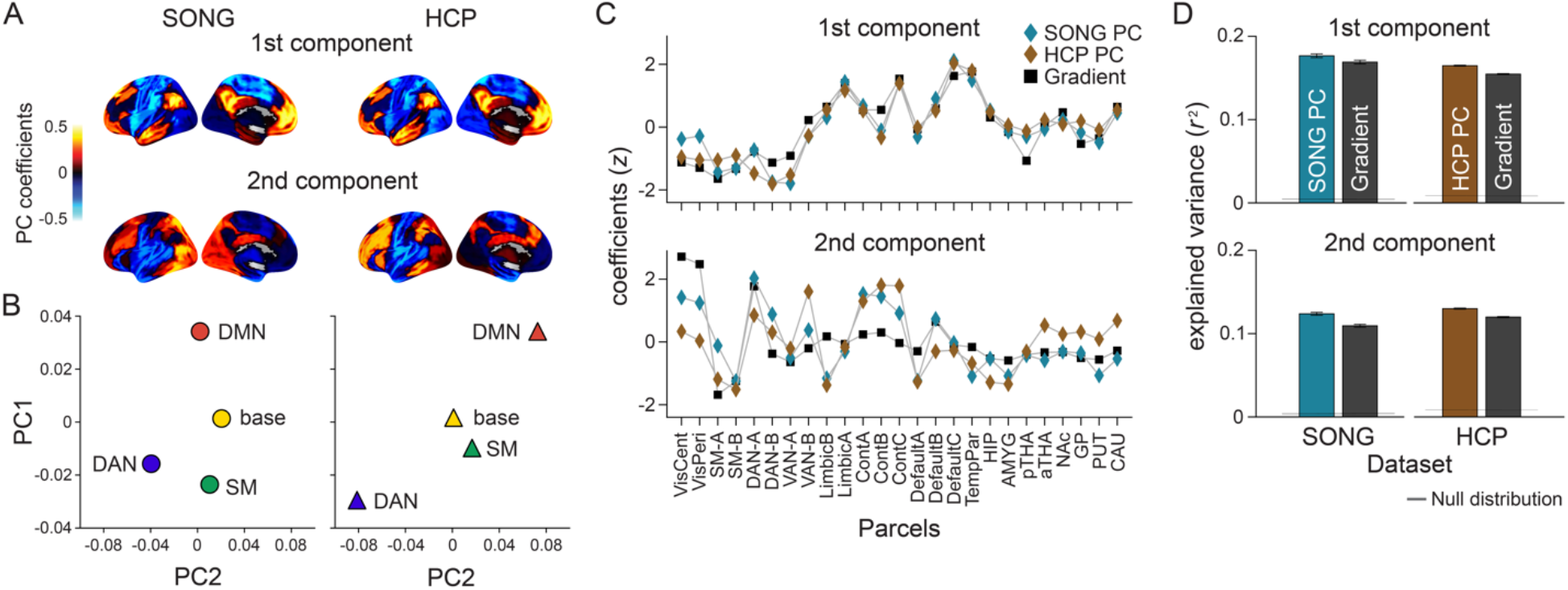
Data-specific principal component analysis on the neural dynamics compared to dynamics summarized along the predefined gradients. **(A)** First and second principal component (PC) coefficients of the SONG (left) and HCP (right) datasets. The lateral and medial views of only left hemisphere are visualized for simplicity. **(B)** Latent neural states of the SONG (left) and HCP (right) datasets situated on the respective PC axes. Positions in the PC space reflect the mean element-wise product of the PC coefficients of the 25 parcels and mean activity patterns of each HMM inferred from the SONG (left) and HCP (right) datasets. **(C)** The first and second gradient values of the 25 parcels (black squares) and the PC coefficients of the SONG (blue diamonds) and HCP (brown diamonds) datasets (*z*-normalized across parcels for visualization). Gray lines connecting the dots are drawn to aid comparisons of the PC and gradient patterns. **(D)** The explained variance of the SONG and HCP network time series from the first and second PCs and gradients. Explained variance is calculated as the mean of squared Pearson’s correlations (*r*^2^) of the 25 parcel time courses and the time courses projected onto the first and second PC or gradient axes. The shaded gray area (which is narrow and thus appears as a horizontal line) indicates the range of the null distribution (mean ± 1.96 × standard deviation), in which the circular-shifted 25 parcel time series were correlated with the PC and gradient time courses. The error bars indicate S.E.M.

**Fig. S8.**
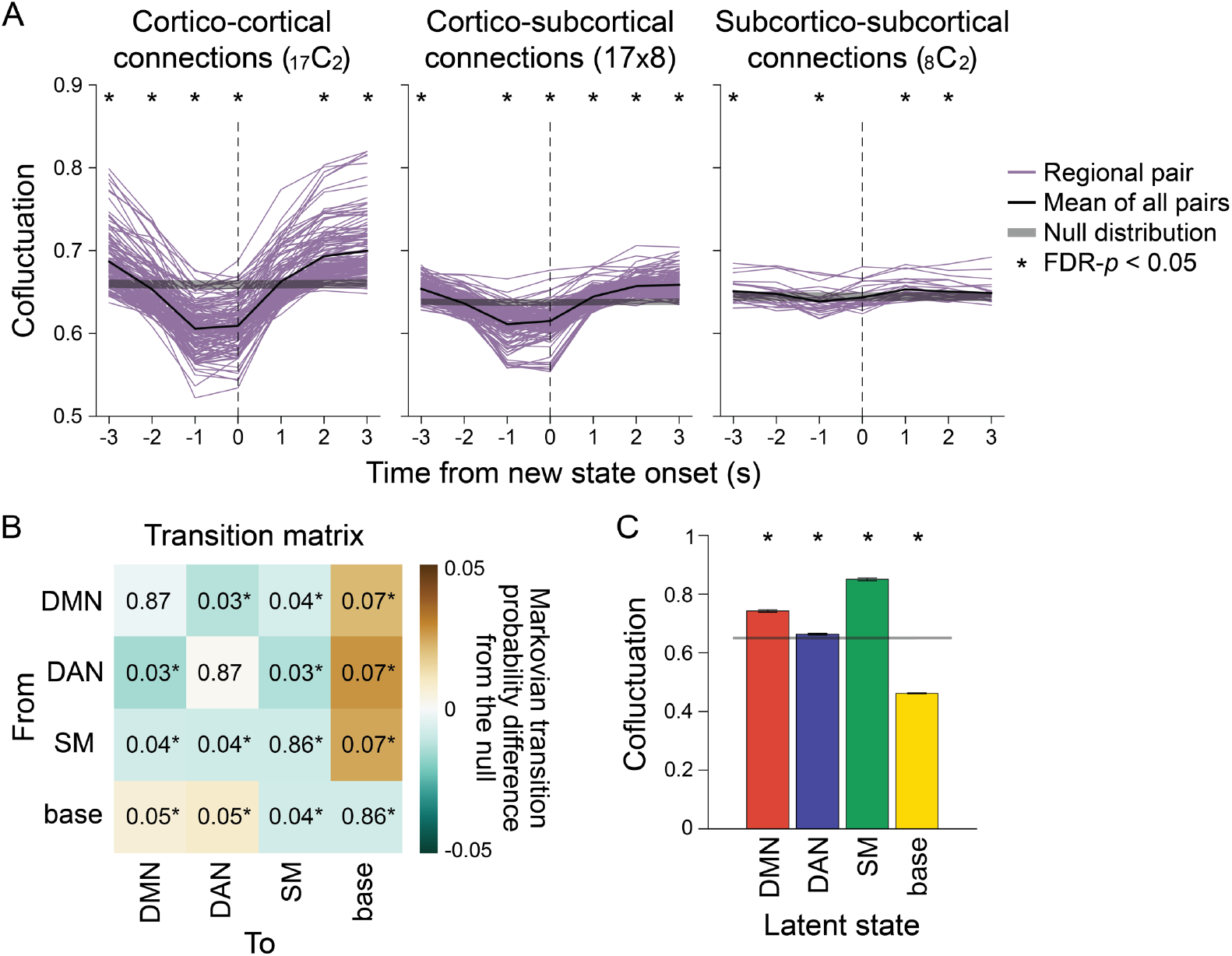
Neural state transitions of the HCP dataset. The figure complements **Fig. 2** which shows the results of the SONG dataset.

**Fig. S9.**
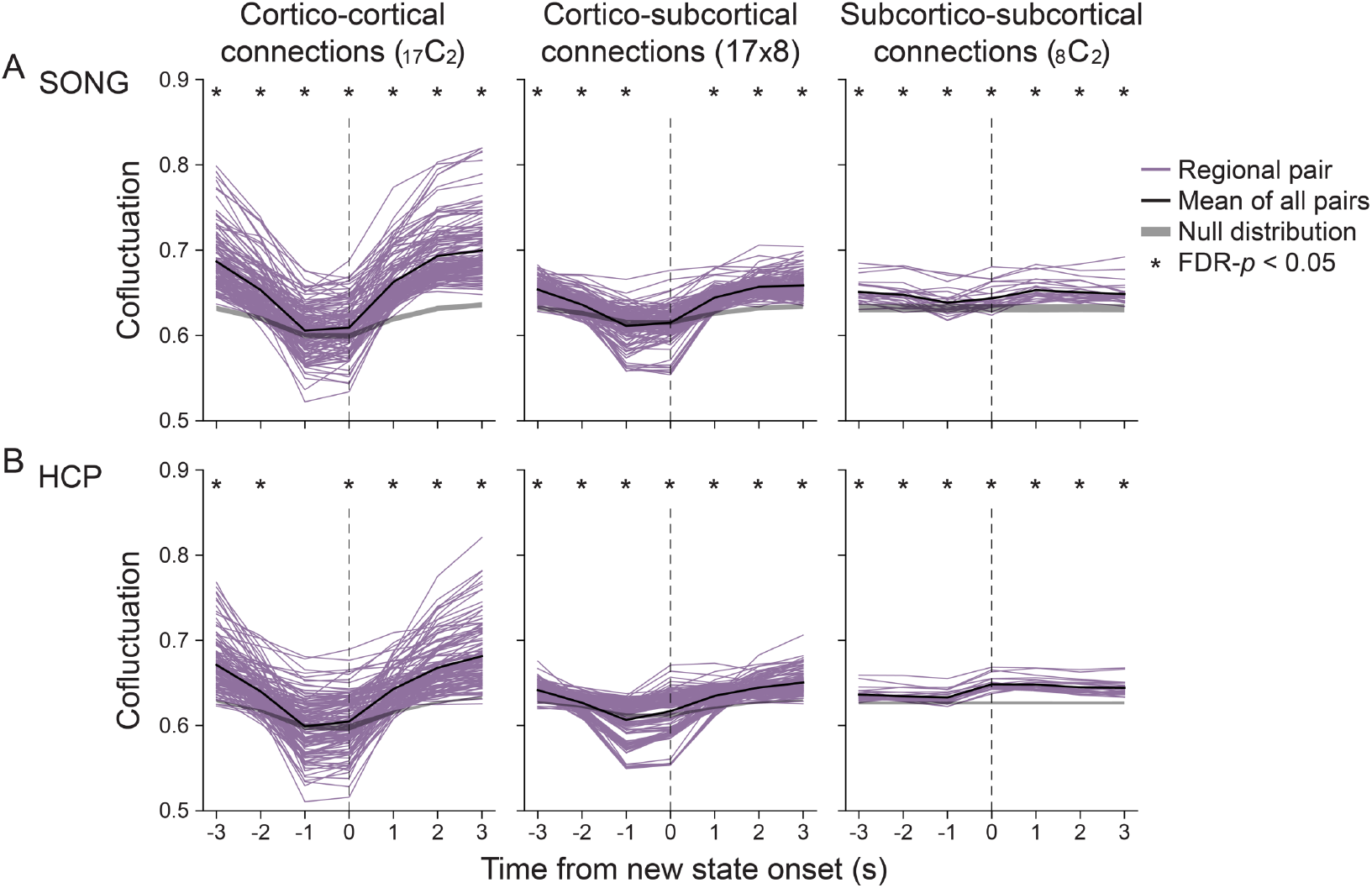
Cofluctuations of all pairs of 25 parcels of the **(A)** SONG and **(B)** HCP datasets, time-aligned to the HMM-derived neural state transitions, compared to a null distribution that was generated differently than **Fig. 2A**. Instead of shuffling the moments of neural state transitions (**Fig. 2A**), we circular-shifted the parcel time series prior to the HMM inference (1,000 iterations, two-tailed non-parametric permutation tests, FDR-corrected for the number of time points). This way of creating null distribution enabled us to ask whether the HMM, by the nature of the model, detects state transitions based on transient decrease in the global synchrony. We observed a decrease in cofluctuation between cortico-cortical and cortico-subcortical regions prior to null state onset (difference between the mean cofluctuation at time *t+3* and *t-1* aligned to new state onset, bootstrapped *p* values < 0.001). However, this decrease was less dramatic than that observed in the actual fMRI time series (two-tailed non-parametric permutation tests, FDR-*p* values < 0.002, corrected for the three pair types). Thus, decreases in cofluctuation prior to state transitions is not simply a byproduct of the computational model.

**Fig. S10.**
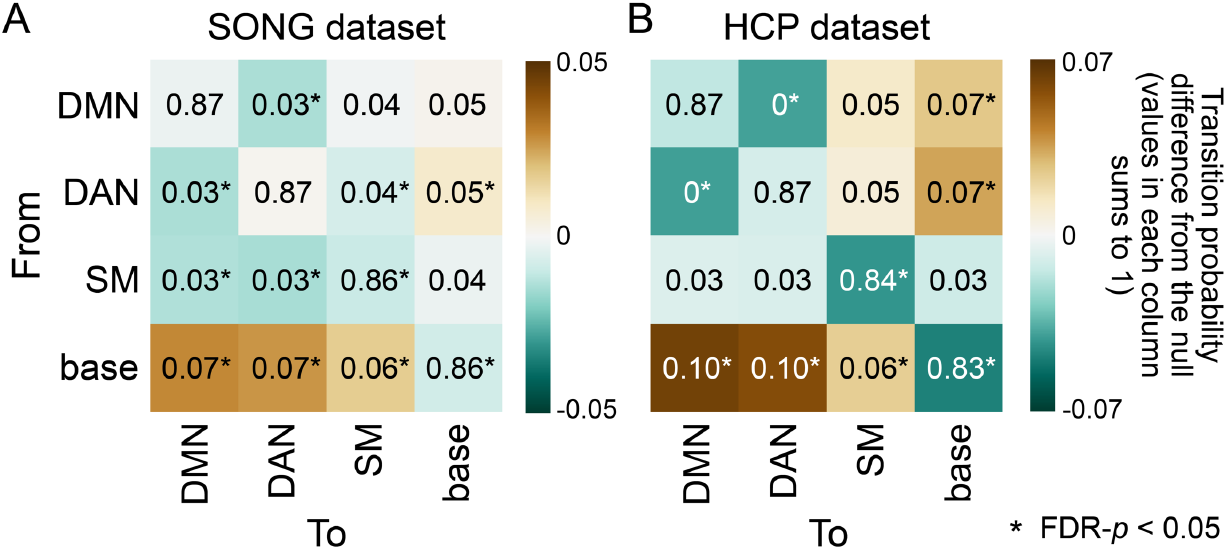
Transition matrix of the **(A)** SONG and **(B)** HCP datasets indicating transition probability from one state (row) to the next (column), such that values in each column sums to 1. Complementing **Fig. 2B** which indicates the probability of transitioning *to* a certain brain state from each of the states (values in each row sums to 1), this figure indicates the probability of transitioning *from* each brain state. The colors indicate differences from the mean of the null distribution where the HMMs were conducted on the circular-shifted time series (asterisks indicate FDR-*p* < 0.05).

**Fig. S11.**
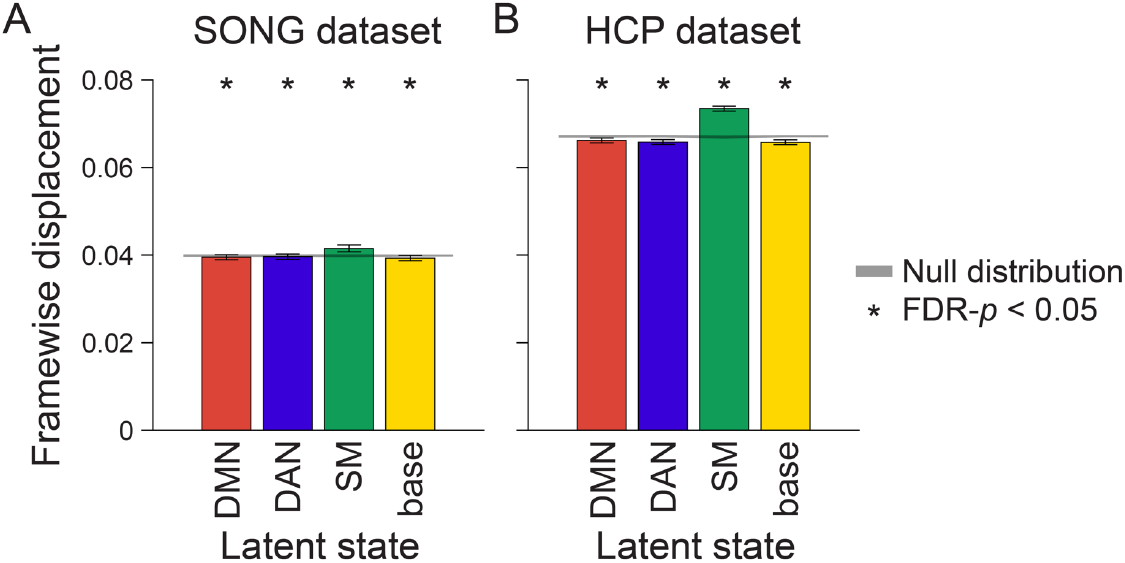
Mean head motion (framewise displacement; FD) at moments of latent neural state occurrence in the **(A)** SONG and **(B)** HCP datasets. FD measured at each time point was averaged within a participant based on the latent state identification, and then averaged across participants. The bar graph indicates the mean of FD from all fMRI runs and participants. The shaded gray area indicates the range of the null distribution (mean ± 1.96 × standard deviation), in which the analyses were conducted on the circular-shifted latent state sequence. FD was measured after motion correction. Higher FD was observed at TRs assigned to the SM state than at TRs assigned to other states (paired *t*-tests, SONG: *t*(187) > 4, HCP: *t*(3091) > 22, both FDR-*p* values < 0.001, corrected for the number of pairwise states), but FD at moments of DMN, DAN, and base states were comparable (SONG: *t*(187) < 1.5, HCP: *t*(3093) < 2, both FDR-*p* values > 0.15).

**Fig. S12.**
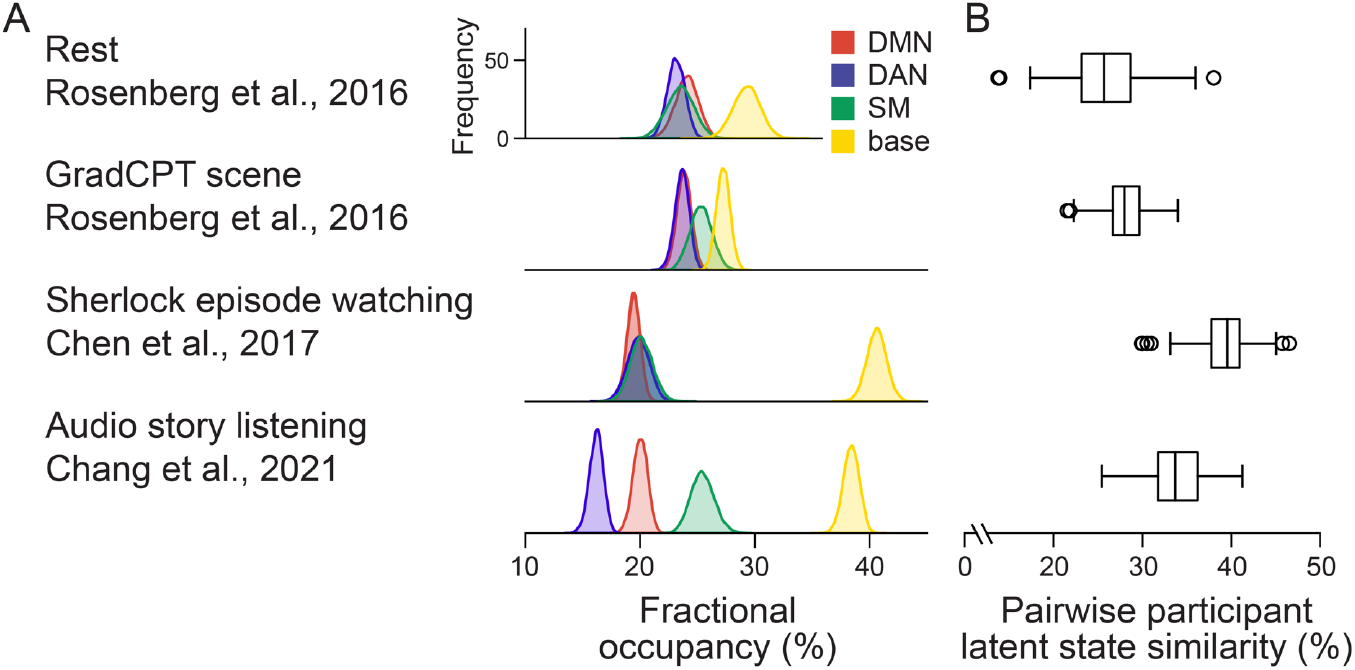
Inferred neural state dynamics of the external datasets from the HMM trained on the SONG dataset. To verify that the inferred neural states are not specific to these datasets, we applied the SONG-trained HMM (i.e., the inferred emission and transition probabilities) to decode latent sequences of the three independent datasets that targeted specific task contexts: the resting-state and gradCPT of Rosenberg et al. (*85*) (N=25), television episode watching of Chen et al. (*64*) (N=16), and story listening of Chang et al. (*66*) (N=25). **(A)** Fractional occupancy of the four latent states per dataset. **(B)** Latent state sequence similarity of all pairs of participants in each dataset. Intersubject synchrony of the latent state sequence was high during television episode watching (*64*) (39.24 ± 3.34%, FDR-*p* < 0.001; comparable to synchrony during the SONG sitcom episodes) and story listening (*66*) (33.84 ± 3.08%, FDR-*p* < 0.001). In comparison, intersubject similarity during rest (*85*) was near chance (25.84 ± 3.79%, though statistically significant FDR-*p* < 0.001). The analysis follows **Fig. 3B-C**.

### Supplementary Text

Participants’ behavioral synchrony closely followed neural synchrony measured with latent state dynamics. To ask whether participant pairs with similar attentional dynamics also exhibited similar neural state dynamics during movie-watching runs, we computed behavioral and neural similarities for all pairs of participants and computed Spearman’s correlations. The behavioral similarity was measured with Pearson’s correlation of either the continuous engagement ratings that each pair of participants completed after the fMRI scans or continuous button responses to gradCPT tasks during the fMRI scans (**Fig. 5A**). The neural similarity was the proportion of time when the two participants exhibited the same latent neural state. The Spearman’s *r* between all pairs’ behavioral and neural similarities were compared with a chance distribution in which the participant indices for behavioral measures were randomly shuffled (FDR-corrected for 3 movie-watching fMRI scan runs).

Here, we further report the degrees to which behavioral attention measures were synchronized across participants. The rated engagement dynamics were highly similar across participants during sitcom episode watching (episode 1: Pearson’s *r* = 0.266 ± 0.179, FDR-*p* = 0.001; episode 2: *r* = 0.287 ± 0.262, FDR-*p* = 0.001; paired comparisons, non-parametric *p* = 0.128). Engagement during documentary watching was synchronized (*r* = 0.155 ± 0.202, FDR-*p* = 0.001) but significantly less so compared to the sitcom episodes (paired comparisons with the two sitcom episodes, both *p* < 0.001), whereas the synchrony of RT variabilities during gradCPT was near chance (face: *r* = 0.062 ± 0.073, FDR-*p* = 0.001; scene: *r* = 0.005 ± 0.053, FDR-*p* = 0.048; paired comparison, *p* < 0.001). These results closely follow the degrees of synchrony in neural state dynamics (**Fig. 3C**).

**Fig. S13.**
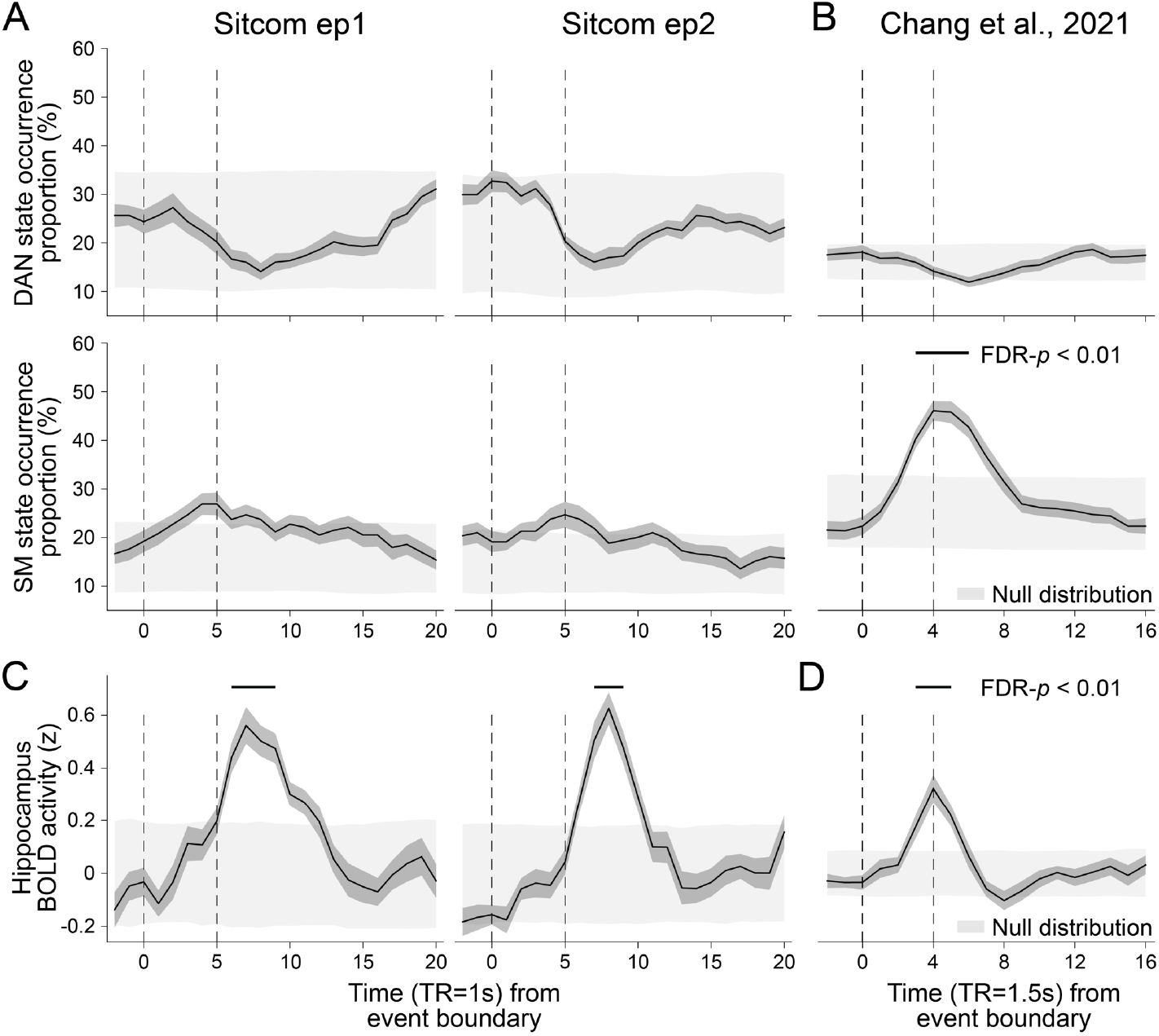
Neural state occurrence and hippocampal BOLD activity time-aligned to narrative event boundaries. **(A-B)** Neural state occurrence after event boundaries. This figure complements **Fig. 4A-B**, which only showed DMN and base state occurrence proportions. Consistent across **(A)** the sitcom episode 1 and 2 runs in the SONG dataset and **(B)** the external dataset, Chang et al. (*66*), no change in the DAN state occurrence was found after event boundaries. No change in the SM state occurrence was found for the sitcom episodes 1 and 2. An increase in SM state occurrence after Chang et al.’s (*66*) event boundaries is likely to be due to ∼6 s of silent pauses in the audio in between the events. Given that the SM state occurred at intermittent rests in between the task blocks (**fig. S15**), the increase in the SM state after event boundaries may be due to a blank period rather than the narrative event change. **(C-D)** BOLD activity of the hippocampal ROI time course (*z*-normalized), time-aligned to narrative event boundaries of the **(C)** SONG dataset’s sitcom episodes and **(D)** Chang et al.’s (*66*) audio story. The hippocampal ROI mask of the MNI template was adopted from Tian et al. (*30*).

**Fig. S14.**
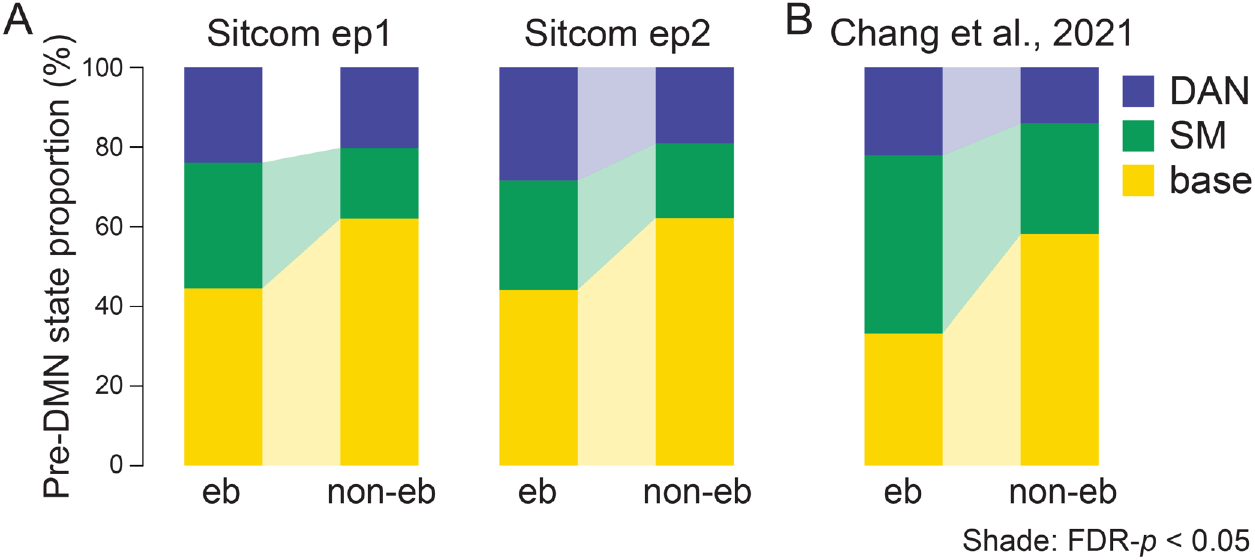
Transitions to the DMN state at narrative event boundaries. This figure complements **Fig. 4C**, which visualizes the schematics of these results. **(A)** The proportion of the DAN, SM, and base states that occurred prior to the DMN state occurrence during the sitcom episodes 1 (left) and 2 (right). Transitions to the DMN state were categorized based on whether the DMN state occurred 5-15 TRs after the event boundaries (event boundary; eb) or not (non-event boundary; non-eb). Paired Wilcoxon signed rank tests were conducted between the eb and non-eb conditions per neural state, and the significant difference was indicated with a colored shade (FDR-*p* < 0.05). **(B)** Same as **A** but in Chang et al. (*66*) dataset. Transitions to the DMN state 4-12 TRs after the event boundaries were compared to those that occurred at the rest of the moments. The proportions were averaged across participants for visualization.

**Fig. S15.**
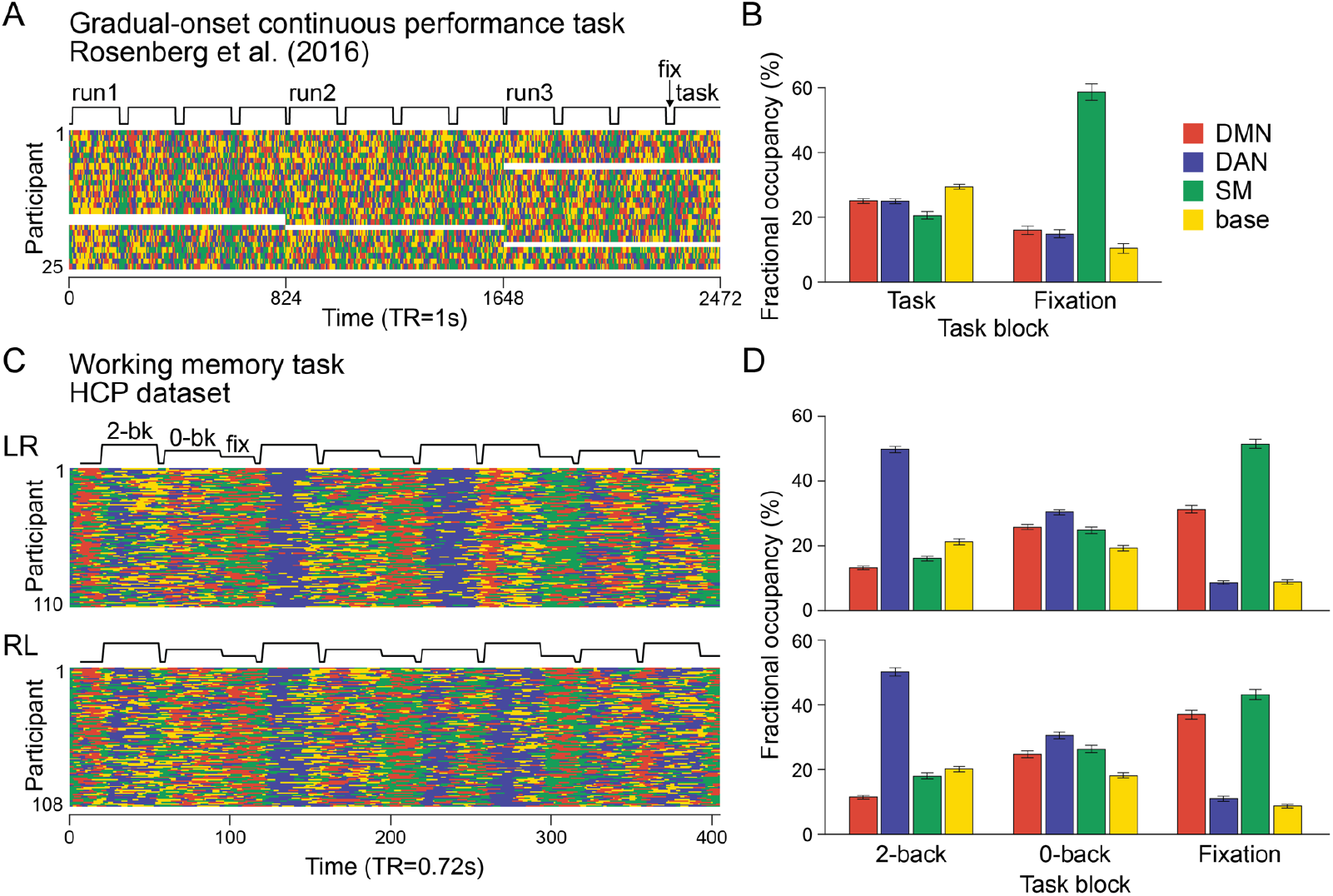
Latent state dynamics during cognitive task blocks, decoded from the HMM trained on the SONG dataset. **(A)** Latent state dynamics of the gradCPT data collected by Rosenberg et al. (*85*) (N=25). The data consists of three runs, with each run consisting of four 3-min task blocks interleaved with 30 s of fixation. A boxcar plot indicates durations of the task and fixation blocks, shifted in time (5 TR; 5 s) to account for hemodynamic response delay. White areas indicate missing fMRI runs. **(B)** Fractional occupancies of the four states were calculated during task and fixation blocks separately and averaged across participants. The bar graph indicates the mean of fractional occupancies calculated per participant. The DMN, DAN, and base states occurred more frequently during task than fixation blocks (paired *t*-tests, *t*(24) > 6, FDR-*p* values < 0.001), whereas the SM state occurred more frequently during fixation blocks (*t*(24) = 12.565, FDR-*p* < 0.001). We observed a significant main effect of neural states (repeated measures ANOVA: F(3,72) = 85.479, *p* < 0.001) and an interaction between neural states and block types (F(3,72) = 119.392, *p* < 0.001). The error bars indicate S.E.M. **(C)** Latent state dynamics of the HCP WM task runs in left-to-right (LR; N=110) and right-to-left (RL; N=108) phase encoding sessions. Each session consists of a single run, with four 2-back WM (25 s), four 0-back WM (25 s), and three fixation blocks (15 s) interleaved, which are indicated in the boxcar plots shifted in time (6 TR; 4.32 s). **(D)** Fractional occupancies of the four states calculated at moments of 2-back WM, 0-back WM, and fixation blocks. In both sessions, DAN state occurrence decreased (paired *t*-tests for all block pairs, LR: *t*(109) > 13, RL: *t*(107) > 10, FDR-*p* values < 0.001), whereas the DMN and SM state occurrence increased (LR: *t*(109) > 3, RL: *t*(107) > 5, FDR-*p* values < 0.001), from 2-back to 0-back to fixation blocks as blocks required less cognitive control and memory load. The base state, on the other hand, occurred comparably during the 2-back and 0-back WM task blocks (LR: *t*(109) = 1.804, FDR-*p* = 0.074, RL: *t*(107) = 1.486, FDR-*p* = 0.140), whereas the it occurred less at fixation blocks (LR: *t*(109) > 10, RL: *t*(107) > 9, FDR-*p* values < 0.001). The repeated measures ANOVA showed the main effect of the neural states (LR: F(3,327) = 117.445, *p* < 0.001; RL: F(3,321) = 126.702, *p* < 0.001) and the interaction effect (LR: F(6,654) = 261.987, *p* < 0.001; RL: F(6,642) = 149.786, *p* < 0.001).

## Footnotes

1 Though our observations align with previous work on the functional roles of the default mode and dorsal attention networks, it is important to keep in mind that the two states are not just characterized by activation of these networks but by patterns of activation and covariation of the whole brain networks. They should be interpreted as “states” rather than isolated functional networks.

2 https://www.color-blindness.com/ishihara-38-plates-cvd-test

3 https://en.wikipedia.org/wiki/High_Kick_Through_the_Roof

4 https://www.youtube.com/watch?v=sL3kLxsy-Lg

5 https://www.youtube.com/c/VIVOTVchannel

6 https://identifiers.org/neurovault.collection:1598

